# Nutrient and prey-sensing signaling pathways converge on the sphingolipid methyltransferase *SMT1* to regulate trap formation in a predatory fungus

**DOI:** 10.64898/2026.09.18.752630

**Authors:** Tsung-Yu Huang, Ching-Ting Yang, Chih-Yen Kuo, A Pedro Gonçalves, Guillermo Vidal-Diez de Ulzurrun, Hillel Schwartz, Yen-Ping Hsueh

**Affiliations:** Molecular and Cell Biology, Taiwan International Graduate Program, Academia Sinica and Graduate Institute of Life Science, National Defense Medical Center, Taipei, Taiwan; Institute of Molecular Biology, Academia Sinica, Taipei 11529, Taiwan; Department of Complex Biological Interactions, Max Planck Institute for Biology, Tübingen, Germany

**Keywords:** Nematode-trapping fungi, Carbon Catabolite Repression Cre1, Pheromone-response MAPK signaling Ste12, Sphingolipid methyltransferase

## Abstract

The nematode trapping fungus *Arthrobotrys oligospora* transitions from saprophytic growth to a predatory lifestyle by forming adhesive traps in response to nutrient limitation and nematode derived cues. How nutrient availability integrates with prey sensing to control trap formation remains unclear. The presence of glucose can suppress trap formation and we found that Cre1, a conserved transcription factor that regulates the carbon catabolite repression pathway in fungi is essential for trap formation. Comparative transcriptomics and functional studies identified *SMT1*, which encodes a sphingolipid C9 methyltransferase, as a Cre1-dependent target that is sufficient to restore trap formation defect in the *cre1* mutant. Smt1 contributes to the formation of sterol-enriched membrane domain at the tip of a growing trap hyphae and thus affects trap morphogenesis. We further discovered that the expression of *SMT1* depends on both Cre1 and another transcription factor Ste12 that acts downstream of the pheromone response MAPK pathway critical for prey-sensing. These findings reveal functional crosstalk between Cre1 and Ste12 where both transcription factors are required for the expression of *SMT1* and demonstrate that a predatory fungus integrates nutrient and prey-derived signals to regulate predatory lifestyle switching.

## Introduction

Fungal development is highly responsive to environmental cues, particularly nutrient availability and host-derived signals, which drive morphological transitions essential for biological processes such as pathogenesis and sexual reproduction. In many fungi, the response to these cues relies on the integration of multiple signaling pathways. For example, in the fission yeast *Schizosaccharomyces pombe* and the red bread mold fungus *Neurospora crassa*, mating requires the convergence of nutrient limitation and activation of the pheromone-response mitogen-activated protein kinase (MAPK) cascade to drive cell polarization and fusion [1, 2]. In the rice blast fungus *Magnaporthe oryzae*, plant-derived hydrophobic surface cues and nutrient starvation cooperatively induce appressorium differentiation through the Pmk1 MAPK and cyclic AMP-protein kinase A (cAMP-PKA) pathways, which regulate transcriptional reprogramming and morphogenesis of this specialized infection structure [3, 4]. Similarly, in the corn smut fungus *Ustilago maydis*, both nitrogen scarcity and host-derived signals contribute to the switch from saprophytic to invasive growth: nitrogen limitation activates the Nit2 transcription factor to regulate nitrogen metabolism and promotes filamentous growth and virulence, while host surface cues activate the Sho1 and Msb2 MAPK cascade to trigger appressorium formation [5, 6]. Understanding how fungi integrate multiple environmental cues to regulate morphological transitions is key to address their adaptive strategies across saprophytic, pathogenic, and predatory lifestyles.

Nematode-trapping fungi (NTF) provide a striking example of morphological plasticity. Under nutrient-depleted conditions, these soil-dwelling predators detect nematode-derived signals and develop specialized traps to capture the nematode prey [7, 8]. In the adhesive-net-forming species such as *Arthrobotrys oligospora* and *A. flagrans*, nematode signals such as ascarosides are sensed by G protein-coupled receptors (GPCRs) that activate MAPK and cAMP-PKA pathways to promote trap formation [9–12]. Nutrient supplementation, including with ammonium sulfate, glucose, or rich media, suppresses trap formation in NTF [13], indicating a convergence between nutrient availability and prey-sensing in regulating fungal predation. While the molecular mechanisms underlying prey sensing in NTF have been gradually elucidated, how nutrient signaling integrates with prey-sensing remains largely unclear.

Carbon catabolite repression (CCR) is a conserved mechanism that prioritizes the utilization of preferred carbon sources, such as glucose, while repressing alternative carbon metabolism [14]. In *S. cerevisiae*, CCR is mediated by the transcription factor Mig1, which represses gluconeogenic genes when glucose is abundant [15]. Glucose uptake activates hexokinase Hxk2, which inhibits the kinase Snf1 and promotes Mig1 nuclear accumulation. Under glucose depletion, Snf1 phosphorylates Mig1, leading to its nuclear export and derepression of target genes [15–17]. In filamentous fungi, the Mig1 homolog Cre1/CreA serves as a central regulator of CCR, and its activity is modulated through phosphorylation and ubiquitin-mediated degradation [18, 19]. Beyond carbon metabolism, Cre1/CreA also influences fungal morphogenesis and pathogenicity. In the plant pathogen *Magnaporthe oryzae*, loss of MoCreA impairs vegetative growth, conidiation, and appressorium formation [20]. In the insect pathogen *Metarhizium robertsii*, loss of MrCre1 reduces conidiation, appressorium formation, and virulence, whereas deletion of *CRE1* in *Beauveria bassiana* causes severe defects in vegetative growth, conidiation, and pathogenicity [21, 22]. Recent work in *Fusarium graminearum* further links nutrient-dependent CreA regulation to the transition from vegetative to invasive growth [23].

Given that glucose suppresses trap formation in *A. oligospora*, we asked whether Cre1 contributes to the regulation of trap formation. The established roles of Cre1 in fungal growth and morphogenesis indicate a potential role for regulating morphological transitions and trap formation in NTF. Here, we investigated how Cre1 contributes to trap formation in response to nematodes and uncovered a regulatory mechanism in which Cre1 and the prey-responsive transcription factor Ste12 converge on *SMT1* to promote trap morphogenesis.

## Results

### Glucose suppresses *A. oligospora* trap formation in a concentration-dependent manner

To investigate how carbon availability regulates trap development, we examined trap formation in *A. oligospora* under varying glucose concentrations. On low-nutrient medium (LNM) without glucose supplementation, *A. oligospora* produced abundant traps after *C. elegans* exposure, whereas increasing glucose concentrations progressively reduced trap numbers, with the strongest suppression at the highest concentration tested, 150 mM (82.6% fewer traps than LNM; Fig 1A and 1B; raw data are available in S2 Table). This suppression was stronger with glucose than with sorbitol at the same concentrations, suggesting that the inhibitory effect is not due solely to osmotic effects (Fig 1A and 1B).

**Fig 1.**
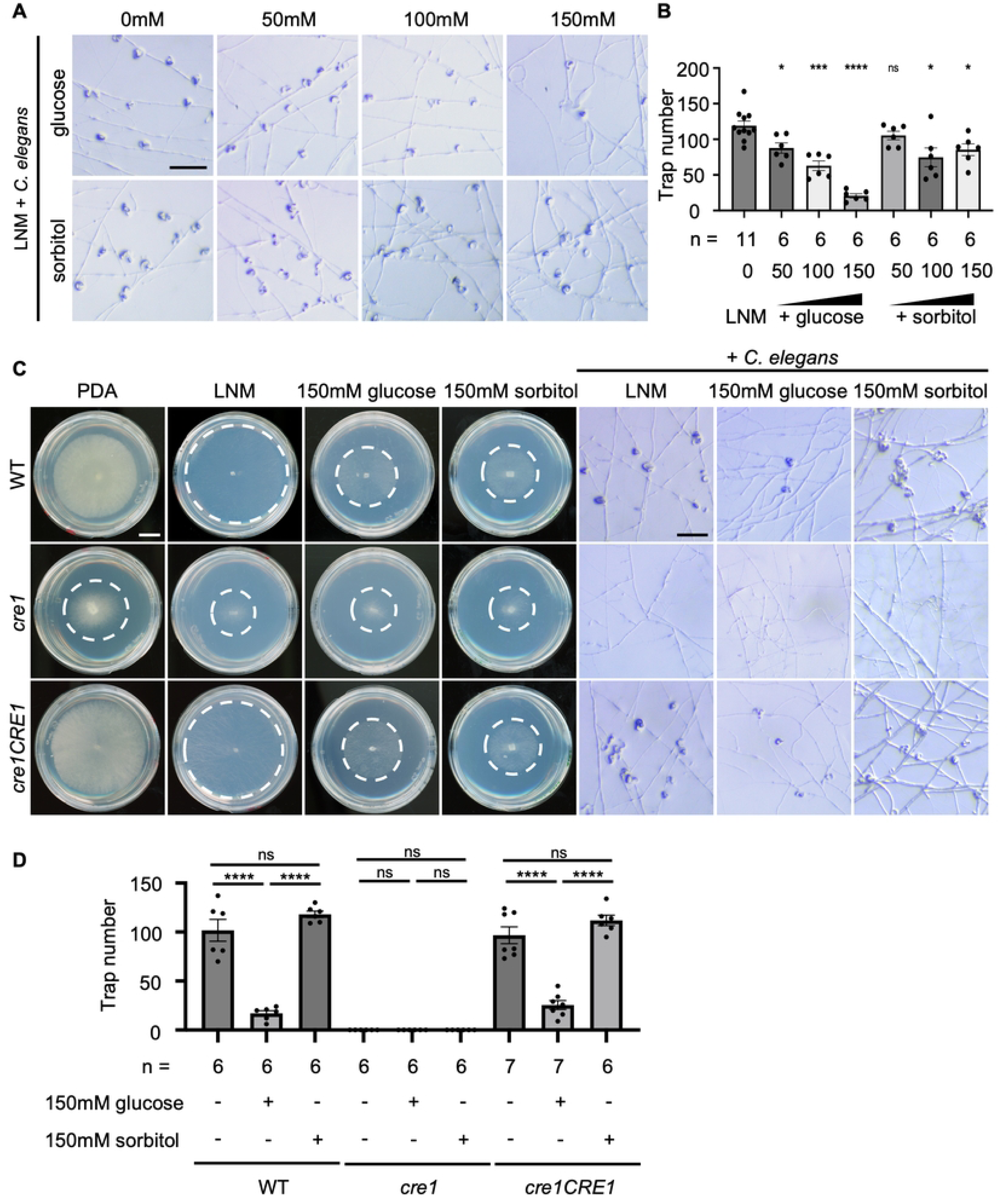
Glucose suppresses trap formation in response to *C. elegans*, and *CRE1* regulates growth and trap formation in *A. oligospora*. (A) Representative brightfield images of traps induced by *C. elegans* in wild type *A. oligospora* grown on LNM plates supplemented with different concentrations of glucose. Sorbitol-supplemented plates were included as an osmolarity control (scale bar = 200 μm). (B) Quantification of trap numbers induced by *C. elegans* in wild type *A. oligospora* grown on LNM plates supplemented with different concentrations of glucose or sorbitol. All statistical comparisons were made relative to the LNM condition. (C) Left: Colony morphology of wild type, *cre1* and *cre1CRE1* strains grown on LNM, LNM supplemented with glucose or sorbitol, or PDA plates for 3 days. White dashed lines indicate colony margins. (Scale bar = 1 cm). Right; Representative brightfield trap images of the same strains on LNM or LNM supplemented with 150mM glucose or sorbitol (Scale bar = 200 μm). (D) Quantification of trap numbers induced by *C. elegans* in wild type, *cre1*, and *cre1CRE1* strains on LNM or LNM supplemented with 150mM glucose or sorbitol. For each strain, statistical comparisons were made between conditions with and without *C. elegans*. Data represent mean ± SEM. Asterisks represent significance levels determined by two-tailed unpaired Student’s t-test; \**P*<0.05, \*\*\**P*<0.001, \*\*\*\**P*< 0.0001, ns: Not significant).

### Cre1 deletion impairs growth and abolishes trap formation in *A. oligospora*

Reasoning that Cre1/CreA homologs mediate carbon catabolite repression pathway and pathogenesis in many fungi, we asked whether *A. oligospora* Cre1 contributes to trap formation. We generated a *cre1* deletion mutant and analyzed its phenotypes. Under both nutrient-rich (PDA) and nutrient-poor (LNM) conditions, the *cre1* mutant formed smaller colonies compared to WT. This slow growth phenotype remains the same when *cre1* mutant was cultured on LNM supplemented with glucose or sorbitol, showing that *cre1* mutant exhibited growth defect (Fig 1C). Furthermore, upon exposure to the nematode *C. elegans*, no traps were formed in the *cre1* mutant under any tested condition, indicating that *CRE1* is essential for trap formation (Fig 1D; raw data are available in S3 Table). Reintroduction of the *CRE1* gene into the *cre1* mutant restored colony morphology, growth, and trap formation (Fig 1C and 1D), indicating that the phenotypes observed were attributed to loss of *CRE1*.

### Glucose and *C. elegans* exposure do not influence Cre1 expression and subcellular localization

To investigate the expression and localization of Cre1, we generated a Cre1-GFP fusion construct driven by the native *CRE1* promoter and introduced into the *cre1* mutant strain. The Cre1-GFP transgene rescued both growth and trap formation in the *cre1* mutant, indicating that the fusion protein is functional (S1A and S1B Fig; raw data are available in S4 Table).

We next examined whether glucose or *C. elegans* exposure influence Cre1 expression and subcellular localization. Through qPCR and western blot analysis, we found that *CRE1* transcript levels and Cre1-GFP protein abundance did not change significantly under either condition (S1C-1E Fig; raw data are available in S5 and S6 Tables). Furthermore, quantitative fluorescent imaging showed that Cre1-GFP localized to nuclei in the hyphae under all conditions tested (S1F Fig).

Sequence alignment of *A. oligospora* Cre1 protein with homologs from *Neurospora crassa*, *Trichoderma reesei*, and *Aspergillus nidulans* revealed strong conservation in the zinc-finger DNA-binding domain and a region spanning residues 292-335 (S2A and S2C Fig) [18, 24]. Structural prediction of *A. oligospora* Cre1 showed a structured zinc-finger domain. In contrast, much of the remaining protein showed low structural confidence, including residues 292-335 (S2B Fig).

To test whether Cre1 is functionally conserved across different ascomycetes, we expressed the *N. crassa CRE1* gene in *A. oligospora cre1* mutant under the native *A. oligospora CRE1* promoter. We found that this restored the defects in growth and trap formation in the *cre1* mutant, indicating that Cre1 function is conserved in two evolutionarily distinct fungi (S2D and S2E Fig; raw data are available in S7 Table).

Given that residues 292-335 of *A. oligospora* Cre1 are highly conserved among fungal Cre1 homologs [18, 25, 26], we tested whether this conserved domain is required for Cre1 function by expressing a truncated Cre1 variant lacking residues 292-335 (Cre1^Δ292–335^) in the *cre1* mutant. Unlike full-length Cre1, *cre1CRE1*^Δ292–335^ variant failed to restore growth or trap formation in *cre1* mutants despite comparable gene expression level (S2D-S2F Fig; raw data are available in S8 Table). These results demonstrate that this conserved domain is critical for the function of Cre1.

### Cre1 regulates the transcriptional response to *C. elegans*

Having found that Cre1 is required for trap formation in *A. oligospora*, we next asked how Cre1 regulates global gene expression in response to *C. elegans*. We performed comparative transcriptomic profiling under four conditions: wild type (WT), WT exposed to *C. elegans*, *cre1* mutant, and *cre1* mutant exposed to *C. elegans* (Fig 2, S9-S12 Tables). Principal component analysis revealed four distinct clusters among the four different conditions, indicating distinct transcriptomes among four conditions (Fig 2A, S9 Table). We first compared the *cre1* mutant to the wild type in the absence of *C. elegans* (*cre1* vs WT, Fig 2B). Differential gene expression (DEG) analysis identified 598 upregulated and 674 downregulated genes in the *cre1* mutant relative to wild type (|beta| ≥ 1.5, where beta is approximately equivalent to log_2_ fold change, FDR-adjusted p < 0.05) (Fig 2B, S10 Table), suggesting that loss of Cre1 causes broad transcriptional regulation. We next examined how the wild type and the *cre1* mutant responded to *C. elegans* (WT+ *C. elegans* vs WT, Fig 2C; and *cre1*+*C. elegans* vs *cre1*, Fig 2D). Wild type strain responds to *C. elegans* exposure by upregulation of 398 genes and downregulation of 546 genes (|beta| ≥ 1.5, FDR-adjusted p < 0.05) (Fig 2C, S11 Table), while the *cre1* mutant showed a weaker transcriptional response, with only 267 genes been upregulated and 214 genes been downregulated (|beta| ≥ 1.5, FDR-adjusted p < 0.05) (Fig 2D, S12 Table).

**Fig 2.**
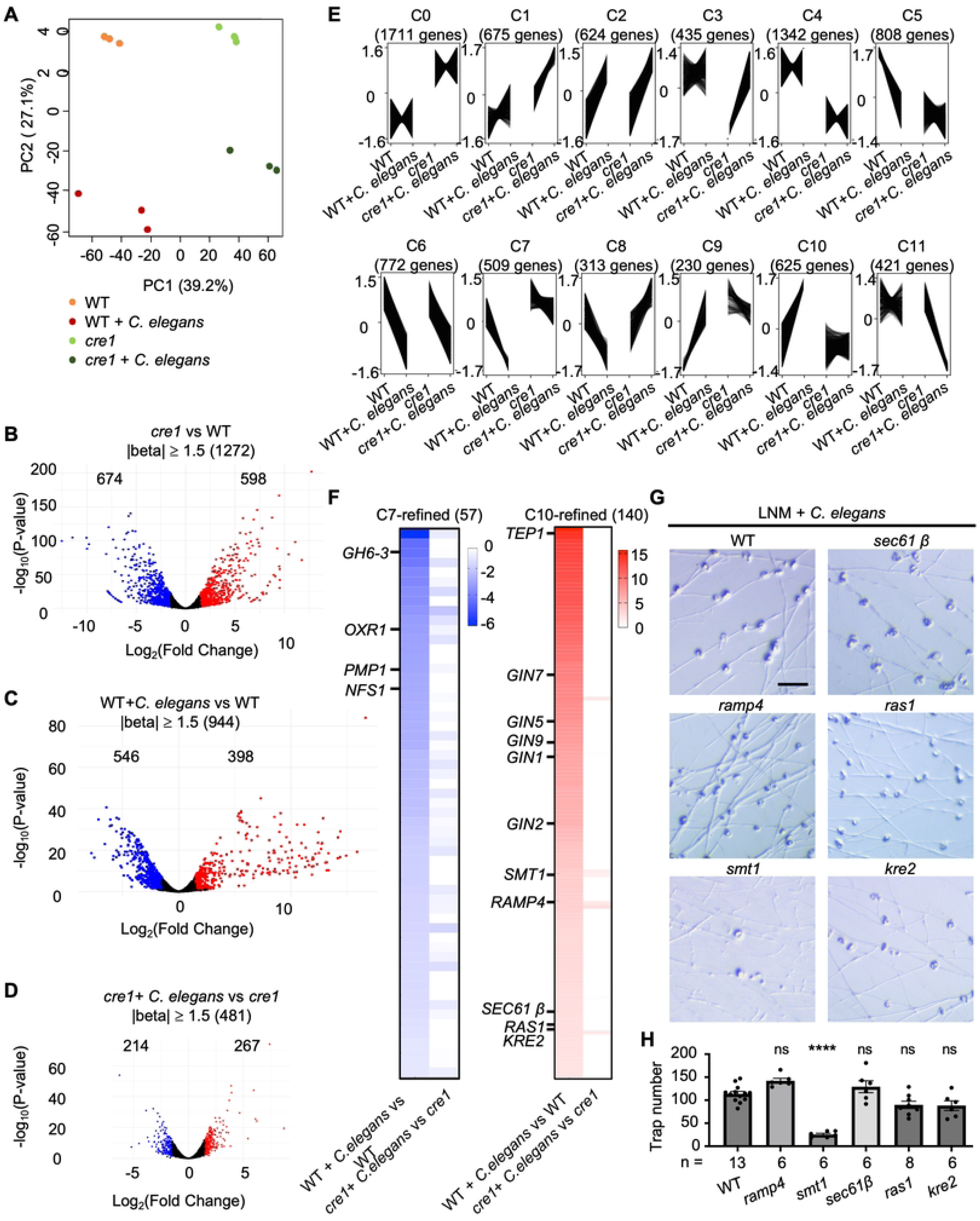
Transcriptomic analysis identifies *SMT1* as a *cre1*-dependent gene involved in trap morphogenesis. (A) Principal component analysis (PCA) of *A. oligospora* transcriptomes from wild-type (WT) and *cre1* mutant strains with or without exposure to *C. elegans*, based on RNA-seq data. Each condition includes three biological replicates, identified by color. (B) Volcano plot displaying differentially expressed genes (DEGs) between WT and *cre1* mutant strains on LNM. DEGs were identified using thresholds of |beta| ≥ 1.5, where beta is approximately equivalent to log_2_ fold change, and FDR-adjusted *P*-value < 0.05. The x-axis represents log₂ fold change (≈ beta) and the y-axis represents -log₁₀(*P*-value). Positively and negatively regulated genes are shown in red and blue, respectively; genes not meeting the DEG thresholds are shown in black. (C) Volcano plot displaying DEGs in WT with or without exposure to *C. elegans*, using the same thresholds and color scheme as in (B). (D) Volcano plot displaying DEGs in the *cre1* mutant with or without exposure to *C. elegans*, using the same thresholds and color scheme as in (B). (E) Cluster analysis of transcript expression profiles identified 12 common expression profiles (C0-C11). Genes were clustered using Clust based on TPM expression profiles across four conditions, with three biological replicates per condition. The x-axis represents the four experimental conditions (WT, WT + *C. elegans*, *cre1*, and *cre1* + *C. elegans*), and the y-axis represents normalized TPM. (F) Heatmaps showing expression levels of 57 DEGs in the high-significance C7-refined gene set and 140 DEGS in the C10-refined gene set in WT and *cre1* mutant strains following exposure to *C. elegans*. Refined gene sets were defined using thresholds of FDR-adjusted *P* < 1×10⁻⁵ and |beta| ≥ 1.5 for the WT + *C. elegans* versus WT comparison. Expression levels are depicted as relative intensity, with more negative beta values shown in darker blues for the C7-refined gene set and higher beta values shown in darker reds for the C10-refined gene set. (G) Representative brightfield images of traps induced by *C. elegan*s in WT and strains carrying deletions of selected genes from the C10-refined gene set (Scale bar = 200 μm). (H) Quantification of trap numbers induced by *C. elegans* in the WT and the indicated gene deletion strains. All statistical comparisons were made relative to WT. Data represent mean ± SEM. Asterisks represent significance levels determined by two-tailed unpaired Student’s t-test; \*\*\*\**P*< 0.0001, ns: Not significant).

To identify potential Cre1-dependent and nematode-responsive genes that might contribute to trap formation, we analyzed transcriptional patterns across all four conditions (WT, WT+ *C. elegans*, *cre1*, and *cre1*+ *C. elegans*). We used the full TPM expression dataset to cluster genes exhibiting similar expression patterns across the four experimental conditions and identified 12 distinct expression profiles (Fig 2E, S13 Table). We focused on clusters C7 and C10, whose expression profiles showed repression (C7) or induction (C10) upon *C. elegans* exposure in the wild type but showed little or no response in the *cre1* mutant (Fig 2E, S13 Table). To define high-confidence Cre1-dependent and nematode-responsive genes within clusters C7 and C10, we filtered genes based on the wild type transcriptional response to *C. elegans* (WT + *C. elegans* vs WT), using thresholds of an FDR-adjusted P-value < 1×10⁻⁵ and |beta| ≥ 1.5. This cutoff was used to prioritize high-confidence DEGs for further analysis. Using these criteria, we refined 509 genes in cluster C7 to 57 genes and 625 genes in cluster C10 to 140 genes (Fig 2F, S14 Table). To distinguish these subsets from the full clusters, we referred to the refined gene sets as C7-refined (57 genes) and C10-refined (140 genes).

Gene Ontology (GO) enrichment analysis of the C7-refined gene set, comprising genes whose repression in response to *C. elegans* was Cre1-dependent, revealed no strongly enriched biological process terms (adjusted P-value=1). However, the “carbohydrate metabolic process” was the most represented term, which may reflect the conserved role of fungal Cre1 homologs in CCR (S3A Fig, S15 Table). We thus prioritized candidates that showed strong differential expression in the wild type compared to in the *cre1* mutant and that had predicted functions possibly relevant to trap formation. We selected four candidates for further study, including the cell wall remodeling glucosyl hydrolase gene *GH6-3*, the oxidative reductase gene *OXR1*, the MARVEL domain transmembrane protein gene *PMP1*, and a nematode-trapping fungi specific gene *NFS1* (Fig 2F, S14 Table). Reasoning that these four genes are strongly down-regulated in the wild type but not in the *cre1* mutant response to *C. elegans*, we overexpressed these genes in the wild type background and examined whether trap formation will be suppressed. However, overexpression of each of the genes in the wild type did not affect trap formation (S3B Fig; raw data are available in S16 Table).

We next conducted GO enrichment analysis of the C10-refined gene set comprising genes strongly upregulated in response to *C. elegans* in the wild type but not in the *cre1* mutant, and similarly failed to identify a strongly correlated term (S3A Fig, S15 Table). We therefore prioritized candidates that were strongly induced by *C. elegans* specifically in the wild type and had predicted functions that might contribute to trap formation (Fig 2F, S14 Table). We excluded predicted secreted proteins, which accounted for 61 of the 140 genes, because based on our previous work, these genes, including the 5 DUF3129 domain containing genes, likely encode proteins enriched in trap cells [27]. We also excluded previously characterized GPCRs (*GIN1*, *GIN2*, *GIN5*, *GIN7*, and *GIN9*) [12] and focused on genes with predicted roles in other cellular processes. We selected five candidates for further study, including the putative sphingolipid C9-methyltransferase gene *SMT1*, the ER stress-associated membrane protein gene *RAMP4*, the protein translocation factor Sec61β, the small GTPase gene *RAS1*, and *KRE2* involved in cell wall remodeling. Deletion of *KRE2, RAS1*, *RAMP4*, or *SEC61β* did not impair trap formation, whereas deletion of *SMT1 strongly* impaired trap formation in response to nematodes (Fig 2G, 2H; raw data are available in S17 Table), demonstrating that *SMT1* is a Cre1-dependent and nematode-responsive gene involved in trap morphogenesis (Fig 2G, 2H).

### *SMT1* functions downstream of both Cre1 and Ste12 during trap formation in *A. oligospora*

Smt1 is a sphingolipid C9-methyltransferase that converts a ceramide intermediate to C9-methylated ceramide during fungal glucosylceramide (GlcCer) biosynthesis [28, 29]. In other fungi, the C9 methylation is critical for growth, membrane integrity, morphogenesis, and virulence [30–33]. *A. oligospora* Smt1 showed high sequence homology with orthologs from *A. nidulans, N. crassa, C. albicans,* and *C. neoformans*, particularly in the predicted transmembrane region and the methyltransferase domain (S4A-S4D Fig) [28]. Deletion of *SMT1* caused severe growth defects, loss of conidiation, and impaired trap morphogenesis, with trap hyphae failing to form proper loops (Fig 3A-3C and S5 Fig; raw data are available in S18 Table). To test whether *SMT1* is a critical downstream gene of Cre1 that contributes to trap formation, we overexpressed *SMT1* in the *cre1* mutant. Indeed, overexpression of *SMT1* in the *cre1* mutant restored trap formation but did not rescue the growth defect of *cre1* (Fig 3A, 3C and S5 Fig). These results suggest that *SMT1* is one of the key downstream targets of *CRE1* required for trap morphogenesis.

**Fig 3.**
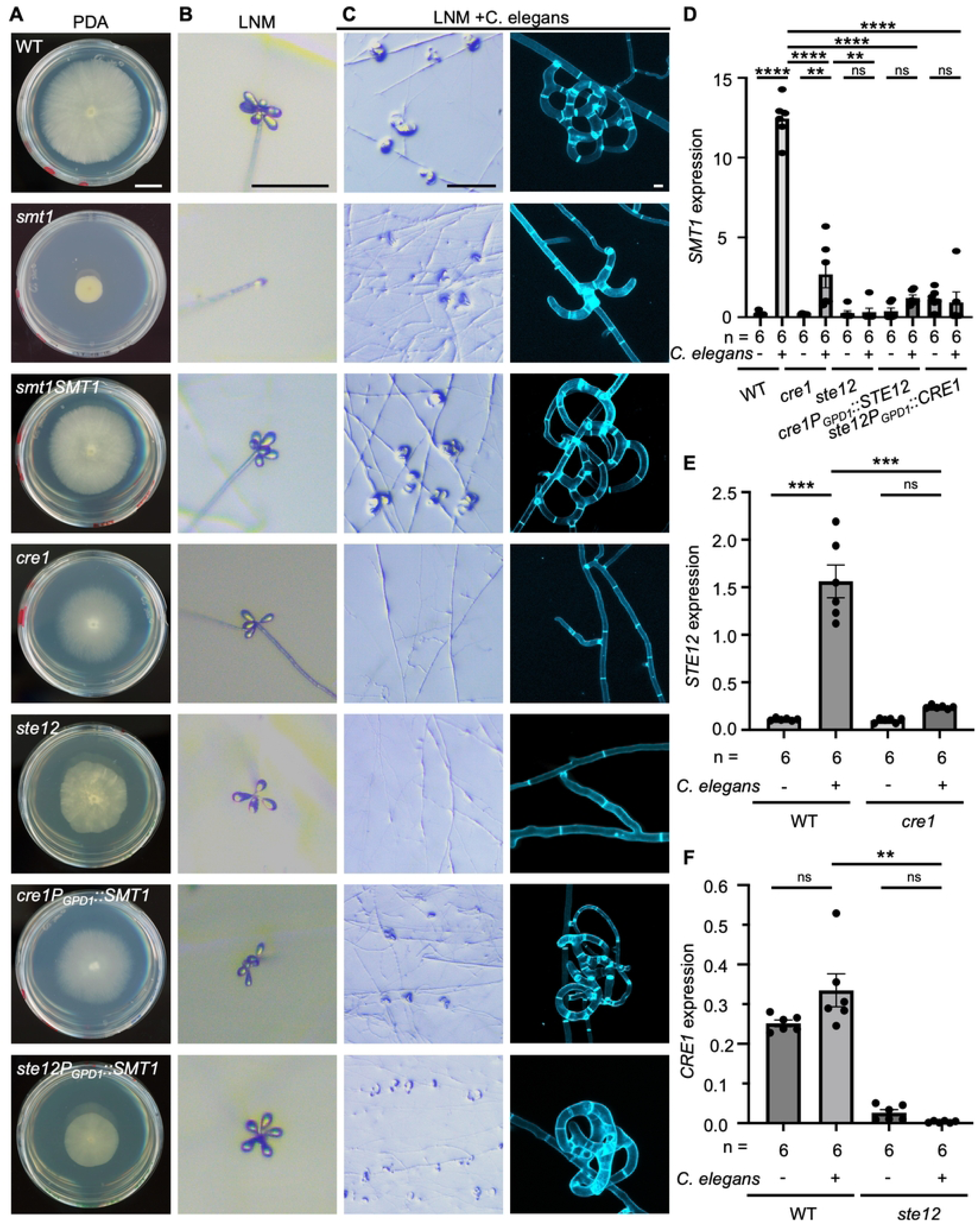
*SMT1* expression depends on both Cre1 and Ste12 and promotes trap formation in *cre1* and *ste12* mutants. (A) Representative images of colony morphology of WT, *smt1*, *smt1SMT1*, *cre1*, *ste12*, *cre1P_GPD1_::SMT1*, and *ste12P_GPD1_::SMT1* grown on PDA plates (scale bar = 1 cm). (B) Representative brightfield images of conidiation in the indicated strains grown on LNM (Scale bar = 200 μm). (C) Left: Representative brightfield images of trap formation in the indicated strains cultured on LNM (Scale bar = 200 μm). Right: Confocal microscopy images of SR2200-stained trap formed by indicated strains after 6h continuous induction with *C. elegans* (Scale bar = 10 µm). For (D-F), the expression levels of *SMT1*, *STE12*, and *CRE1* were evaluated by qPCR in the indicated strains with or without exposure to *C. elegans*. *GPD1* was used as the normalization control. (D) *SMT1* expression was evaluated in WT, *cre1*, *ste12*, *cre1P_GPD1_::STE12*, and *ste12P_GPD1_::CRE1* strains. (E) *STE12* expression was evaluated in WT and *cre1* strains. (F) *CRE1* expression was evaluated in WT and *ste12* strains. Data represent mean ± SEM. Asterisks represent significance levels determined by two-tailed unpaired Student’s t-test; \*\**P*<0.01, \*\*\**P*<0.001, \*\*\*\**P*< 0.0001, ns: Not significant.

Given that Ste12 mediates prey-sensing MAPK signaling and is required for trap formation in *A. oligospora* [9], we asked whether *SMT1* expression also depends on Ste12. We observed that *SMT1* was strongly induced by *C. elegans* in the wild type but not in the *ste12* mutant, suggesting that *SMT1* is a Ste12-dependent gene upon exposure to nematodes (Fig 3D; raw data are available in S19 Table). Overexpression of *SMT1* in the *ste12* mutant was sufficient to restore trap formation but did not rescue the growth defect. These results suggest that *SMT1* is also one of the key downstream targets of *STE12* required for trap morphogenesis (Fig 3A, 3C and S5 Fig).

### *SMT1* induction depends on both Cre1 and Ste12

To further examine how *CRE1* and *STE12* function within the regulatory network regulating *SMT1* expression and trap development, we analyzed their expression in reciprocal mutants. We found that *CRE1* and *STE12* each promoted the expression of the other gene (Fig 3E and 3F). Induction of *STE12* expression in response to *C. elegans* was strongly reduced in the *cre1* mutant compared with the wild type (Fig 3E; raw data are available in S20 Table). *CRE1* expression, which did not change in response to *C. elegans* in the wild type, was nearly abolished in the *ste12* mutant both in the presence or absence of *C. elegans* (Fig 3F; raw data are available in S21 Table). We next conducted reciprocal overexpression experiments using a constitutively active promoter (S6 Fig; raw data are available in S22 and S23 Tables). In both the *cre1* and *ste12* mutants, overexpression of the other transcription factor failed to restore the trap formation defects (S6C Fig). Furthermore, *SMT1* expression remained very low in both strains, indicating that both Cre1 and Ste12 are required for *SMT1* induction in response to *C. elegans* (Fig 3D).

### Smt1 is required for sterol enrichment at hyphal tips during trap morphogenesis

To examine subcellular localization of Smt1 during vegetative hyphae growth and trap development, Smt1 was C-terminally tagged with GFP under its endogenous promoter. The Smt1-GFP fusion restored growth and trap formation in the *smt1* mutant, indicating that the Smt1-GFP retained Smt1 function (S7 Fig; raw data are available in S24 Table). During vegetative hyphae growth, Smt1-GFP was distributed throughout the cell and concentrated near hyphal tips (Fig 4A). In trap hyphae, Smt1-GFP preferentially localized to the inner rim and hyphal tips (Fig 4A and 4C; raw data are available in S25 Table). To quantify this spatial distribution, we compared mean fluorescence intensity between the two sides of vegetative hyphae and trap hyphae (Fig 4C, S25 Table). Smt1-GFP fluorescence was similarly distributed between the two sides of vegetative hyphae but was 1.2-fold higher along the inner rim than the outer rim of trap loops (Fig 4C, S25 Table). These findings suggest that asymmetric Smt1 localization may contribute to membrane remodeling at the inner rim of the trap loop during trap morphogenesis.

**Fig 4.**
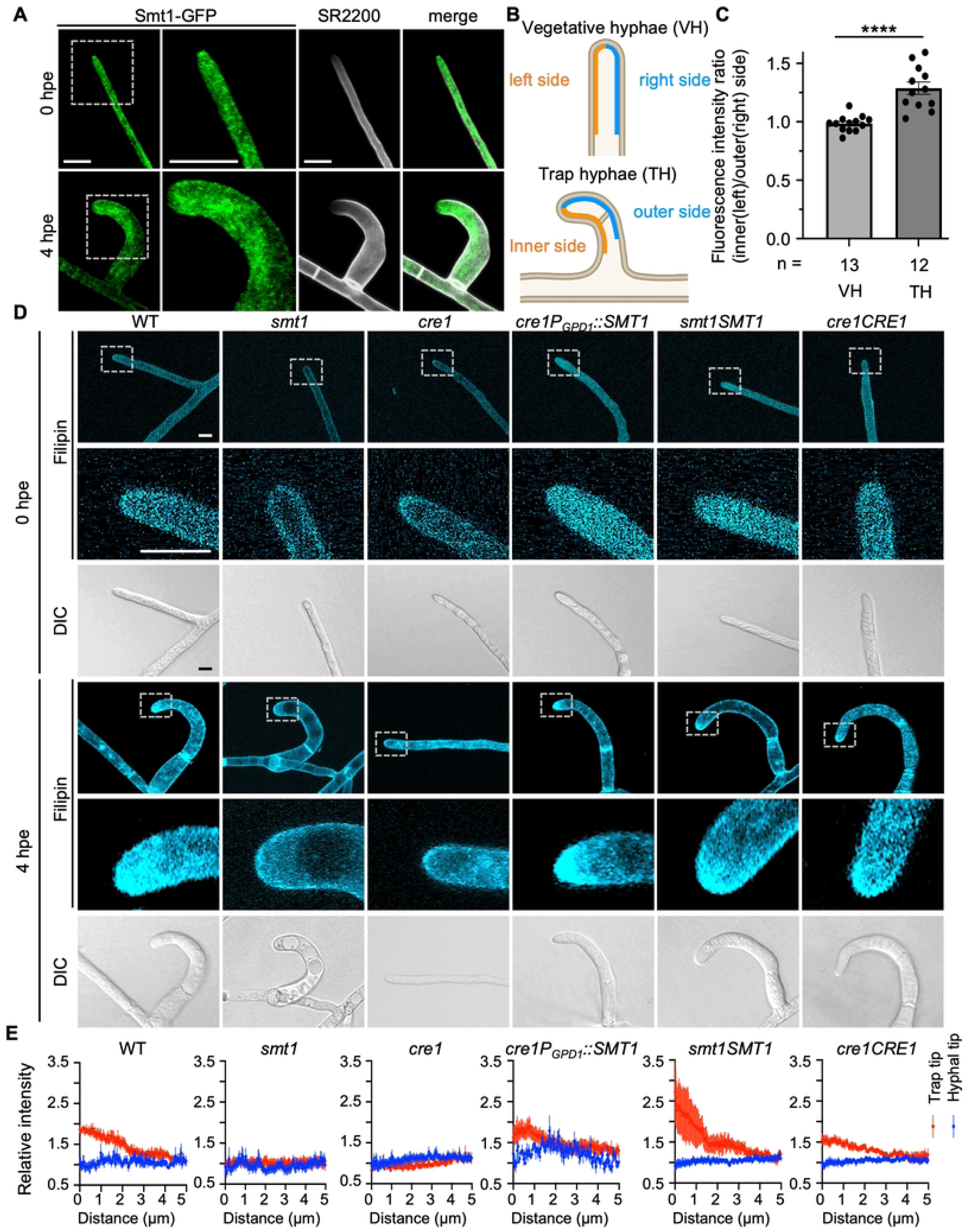
*SMT1* is required for sterol enrichment at the tips of developing trap hyphae. (A) High-resolution confocal images of vegetative hyphae and traps formed by the *smt1SMT1-GFP* strains before and 4h after exposure to *C. elegans*. Fungal cell walls were stained with SR2200. Enlarged regions are indicated by white dashed rectangles in the lower-magnification images (Scale bar = 10 µm). (B) Schematic illustrating the regions used for Smt1-GFP fluorescence quantification in vegetative hyphae (VH) before exposure to *C. elegans* and in trap hyphae (TH) 4h after *C. elegans* exposure in the *smt1SMT1-GFP* strain. (C) Relative Smt1-GFP fluorescence intensity ratios measured between the left and right sides of vegetative hyphae (VH) before *C. elegans* exposure and between the inner and outer sides of trap hyphae (TH) 4 h after *C. elegans* exposure in *smt1SMT1-GFP* strain. The mean fluorescence intensity was calculated for each side, and the left-to-right or inner-to-outer fluorescence ratio was used for quantification, respectively. Statistical comparisons were performed between VH and TH. Data represent mean ± SEM. Asterisks represent significance levels determined by two-tailed unpaired Student’s t-test; \*\*\*\**P*< 0.0001. (D) High-resolution confocal images of hyphae and traps formed by WT, *smt1*, *cre1*, *cre1P_GPD1_::SMT1*, *smt1SMT1*, and *cre1CRE1* strains with or without *C. elegans* exposure. Samples were stained with filipin to visualize sterol distribution. Enlarged regions are indicated by white dashed rectangles in the lower-magnification images. (Scale bar = 5 µm). (E) Relative filipin fluorescence intensity profiles measured along the apical region of trap and vegetative hyphal tips in WT, *smt1*, *smt1SMT1*, *cre1*, *cre1P_GPD1_::SMT1*, and *cre1CRE1* strains grown on LNM with or without exposure to *C. elegans*. Given that the *cre1* mutant did not form traps, measurements were taken from branching hyphal tip. Fluorescence was quantified within a rectangular ROI (5 µm in length × 1.9 µm in width) extending from the hyphal tips. The x-axis indicates distance from the hyphal tip and the y-axis indicates relative fluorescence intensity calculated after subtracting fluorescence measured from a randomly selected region of vegetative hyphae using the same ROI size. Data points represent mean relative fluorescence intensity at each distance, with error bars indicating ± SEM. Six hyphal tips were analyzed in each condition.

Sphingolipid and ergosterol are components of specialized membrane domains known as lipid rafts, which contribute to membrane organization and cell polarity in fungi [29, 31, 34, 35]. We therefore asked whether membrane lipid organization changes during trap morphogenesis and whether Smt1 contributes to this process. To address this, we used filipin, a sterol-binding fluorescent dye [36], to examine membrane sterol distribution in wild type, *smt1*, *cre1*, *cre1P_GPD1_::SMT1*, *smt1SMT1*, and *cre1CRE1* in the presence or absence of *C. elegans*. In the absence of *C. elegans*, filipin fluorescence was distributed throughout vegetative hyphae without detectable enrichment at vegetative hyphal tips (Fig 4D and 4E; raw data are available in S26 Table). Following *C. elegans* exposure, filipin fluorescence became strongly enriched at the tips of developing trap hyphae in the wild type strain, with quantitative analysis showing significantly higher fluorescence at trap hyphal tips than at vegetative hyphal tips (Fig 4D and 4E, S26 Table). The *smt1SMT1*, *cre1P_GPD1_::SMT1*, and *cre1CRE1* strains also showed increased filipin fluorescence at the tips of developing trap hyphae (Fig 4D and 4E, S26 Table). In contrast, the *smt1* mutant formed morphologically defective traps but showed no significant increase in filipin fluorescence at trap hyphal tips compared with vegetative hyphal tips (Fig 4E, S26 Table), suggesting that *SMT1* is required for the formation of sterol-rich membrane at trap hyphal tips during trap development. Since the *cre1* mutant failed to form traps, filipin fluorescence was quantified at branching hyphal tips under trap-inducing conditions, and no filipin enrichment was detected compared with vegetative hyphae (Fig 4D and 4E, S26 Table). Together, these results indicate that the formation of sterol-rich membrane at developing trap hyphal tips is a feature of trap morphogenesis and that Smt1 contributes to establishing this membrane organization.

## Discussion

Trap formation in *A. oligospora* requires integration of environmental information associated with nutrient availability and nematode prey [10, 27]. Our findings support a model in which these distinct inputs converge on *SMT1*, placing Smt1 downstream of both the carbon-responsive regulator Cre1 and the prey-responsive regulator Ste12. Although Cre1 homologs are primarily known for mediating carbon catabolite repression in other fungi, our results indicate that Cre1 in *A. oligospora* also contributes to trap development by inducing *SMT1* expression. Likewise, Ste12, which promotes trap formation in response to prey signals [9], contributes to *SMT1* induction. The inability of Cre1 and Ste12 to substitute for one another, together with the partial rescue of trap formation in both mutants by *SMT1* overexpression, suggests that the two transcription factors provide distinct regulatory inputs that converge on *SMT1* rather than acting in a simple linear pathway. This regulatory arrangement may allow *A. oligospora* to integrate nutrient status and prey detection to coordinate trap morphogenesis.

Our findings indicate that Cre1 contributes to trap morphogenesis in *A. oligospora*, which extends beyond its canonical function in carbon catabolite repression that has been well characterized [18, 37]. In several fungal pathogens, Cre1 also contributes to the development of infection-related structures and regulates broader developmental processes, including vegetative growth, conidiation, and germination [20, 21]. Thus, the developmental role of Cre1 in *A. oligospora* is consistent with the broader functions of Cre1 observed in other fungal pathogens, while the mechanism by which glucose status influences Cre1-dependent trap formation remains to be further explored in the future. Cre1 regulation by carbon availability varies among fungi, with changes in transcript abundance reported in *T. reesei* and changes in nuclear localization reported in *B. bassiana and F. graminearum* [22, 23, 38]. In *A. oligospora*, however, neither Cre1 abundance nor nuclear localization was altered by glucose or *C. elegans* exposure, suggesting that Cre1 may be regulated differently during trap development. Despite these differences, Cre1 function is conserved, as demonstrated by complementation with the *N. crassa* Cre1 homolog and by the requirement for the conserved Cre1 292-335 region [18, 26], in trap development. Thus, *A. oligospora* retains certain conserved Cre1 functions while incorporating it into the regulatory network controlling predatory morphogenesis.

Ste12 provides another transcriptional input into trap development through the pheromone responsive MAPK pathway [9]. Ste12-family transcription factors are conserved regulators of fungal developmental transitions and function downstream of pheromone-response MAPK signaling in processes such as mating, filamentous growth, and morphogenesis [2, 39–41]. In *A. oligospora*, Ste12 promotes trap formation in response to prey signals [9], and our findings further identify *SMT1* as a Ste12-dependent gene required for trap morphogenesis. This finding extends the role of Ste12-associated transcriptional regulation to the control of apical membrane organization during trap development in *A. oligospora*.

The convergence of Cre1- and Ste12-dependent regulation *on SMT1* may represent one point at which multiple signaling pathways are integrated during trap development. Previous studies in filamentous fungi have shown that MAPK- and cAMP-PKA signaling can influence Cre1/CreA regulation and enable pathway crosstalk during morphological differentiation [23, 42, 43]. In *A. nidulans*, pheromone-response MAPK components interact with HOG pathway kinases to regulate CreA localization [42]. In *N. crassa*, disruption of the PKA regulatory subunit alters *CRE1* expression and causes polarity defects, whereas *CRE1* deletion partially suppresses these defects and reduces PKA activity [43]. In *F. graminearum*, cAMP-PKA activation promotes nuclear retention of CreA, linking nutrient-responsive signaling to CreA regulation during fungal developmental transitions [23]. In *A. oligospora*, trap morphogenesis is regulated by multiple signaling pathways, including the pheromone-response MAPK [9], HOG [44], cell wall integrity [45], and cAMP-PKA pathway [10]. Although these pathways have not been shown to converge directly on *SMT1*, our finding that *SMT1* expression depends on both Cre1 and Ste12 raises the possibility that Smt1 serves as an integration point for upstream signaling during trap development.

Trap morphogenesis involves coordinated reorientation of hyphal growth and localized bending, accompanied by the recruitment of polarity machinery to the trap apex [46]. Polarized growth in fungi depends on ergosterol- and sphingolipid-rich membrane microdomains, with C9-methylation of the sphingolipid GlcCer affecting membrane organization and sterol-rich apical domains contributing to the localization of polarity factors [29, 31, 47, 48]. In fungi, septin localization is influenced by membrane curvature and interactions with phosphoinositides and sphingolipids [49, 50], suggesting that membrane lipid organization may contribute to the asymmetric recruitment of septins during trap morphogenesis. This relationship is particularly relevant to Smt1, a sphingolipid biosynthesis enzyme that is also asymmetrically enriched at the inner rim of developing traps. Our previous study has revealed that septins are asymmetrically localized and enriched at the inner rim of developing traps [46]. In this work, we further demonstrate that Smt1 also exhibits an asymmetrical localization pattern in the trap cells and contributes to the spatial organization of membrane sterols during trap morphogenesis. Whether Smt1-dependent membrane organization contributes to the asymmetric recruitment of septins at the trap inner rim remains to be determined.

In conclusion, our findings support a model in which nutrient- and prey-responsive signals converge on *SMT1* to regulate trap morphogenesis in *A. oligospora* (Fig 5)*. SMT1*, a sphingolipid C9-methyltransferase regulated by both the carbon-responsive regulator Cre1 and the prey-responsive transcription factor Ste12, is required for the formation of sterol-rich membrane at the tips of developing trap hyphae. The inability of Cre1 and Ste12 to compensate for one another further indicates that these transcription factors make distinct contributions to trap formation. This model links Cre1-Ste12 transcriptional regulation with Smt1-dependent sterol organization that is critical for the formation of traps and enables the transition to a predatory lifestyle in a nematode-trapping fungus.

**Fig 5.**
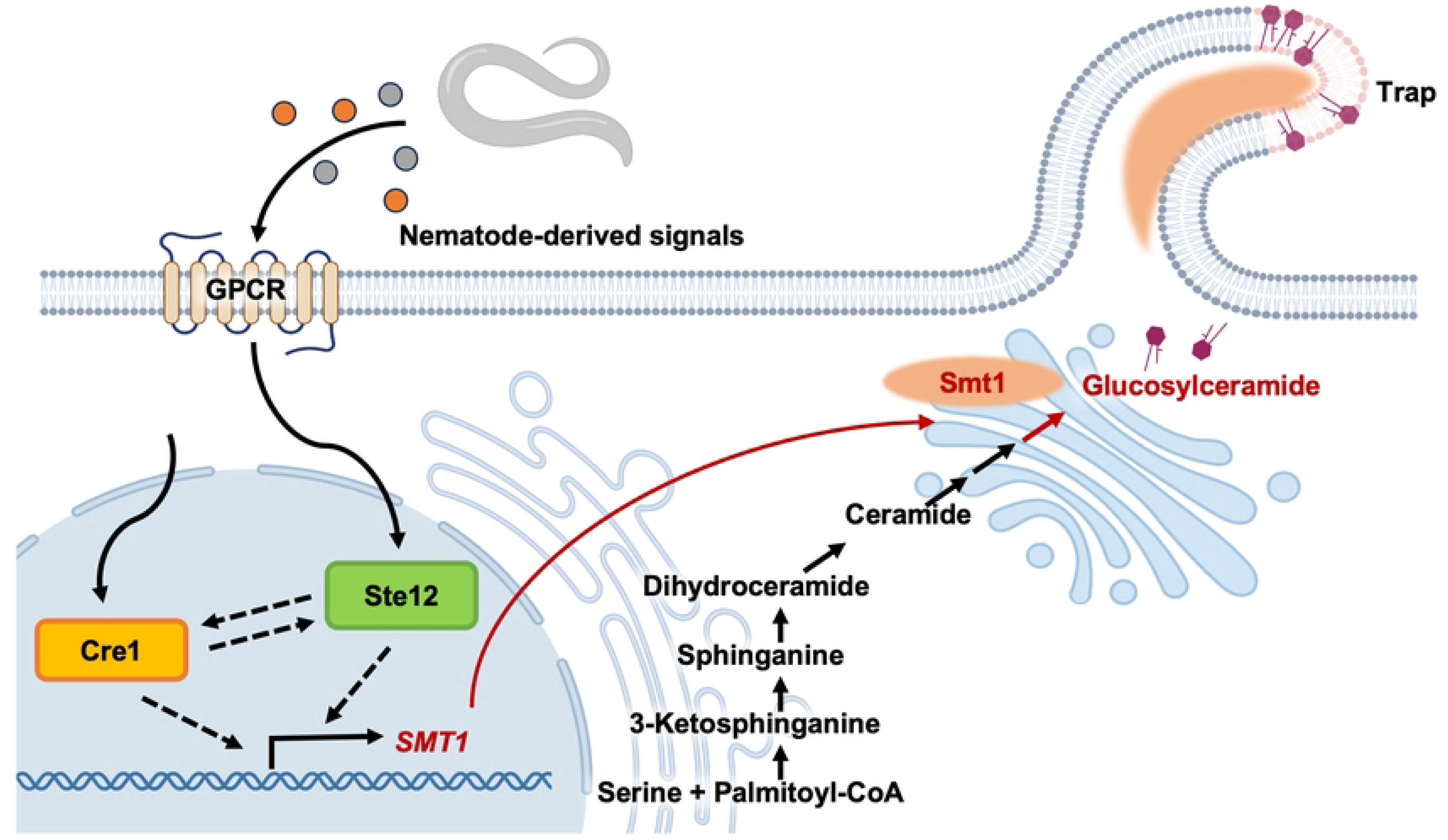
Schematic model of Cre1- and Ste12-dependent regulation of *SMT1* during trap development. Nematode-derived signals activate the pheromone-response pathway and its downstream transcription factor Ste12, while Cre1 provides a distinct regulatory input during trap development. Both Cre1 and Ste12 are required for trap formation and converge to promote *SMT1* expression. Smt1 is predicted to function as a sphingolipid C9-methyltransferase during fungal glucosylceramide biosynthesis and contributes to sterol enrichment at the tips of developing trap hyphae, potentially through effects on sphingolipid composition and membrane organization. Together, Cre1- and Ste12-dependent regulation of *SMT1* and Smt1-associated membrane organization contribute to trap morphogenesis during the developmental transition to fungal predation.

## Materials and methods

### Strains and culture conditions

*Arthrobotrys oligospora* strains used in the study are listed in S1 Table. Cultures were routinely maintained on potato dextrose agar (PDA; Difco) as previously described [9]. For protoplast preparation, liquid cultures were grown in potato dextrose broth (PDB; Difco) following established methods [9]. Trap induction assays were conducted on low-nutrient medium (LNM) agar, composed of 2% agar, 1.66 μM MgSO₄, 5.4 μM ZnSO₄, 2.6 μM MnSO₄, 18.5 μM FeCl₃, 13.4 μM KCl, 0.34 μM biotin, and 0.75 μM thiamin, optionally supplemented with glucose as described previously [9]. *Caenorhabditis elegans* N2 strains were maintained on nematode growth medium (NGM) seeded with *Escherichia coli* OP50 as a food source according to standard methods [51]. All fungal cultures were incubated at 25°C.

### Determination of minimal glucose concentration for trap suppression

Trap induction assays and trap quantification in response to *C. elegans* were performed as described previously [52]. Wild type *A. oligospora* (TWF154) was cultured on 3.5 cm LNM agar Petri plates at 25°C for 48 h. 30 *C. elegans* N2 animals at the young adult stage were added to each fungal plate and incubated for 6 h. For experiments involving *cre1P_GPD1_::STE12* and *ste12P_GPD1_::CRE1* strains, fungal cultures were exposed to around 200 *C. elegans* N2 animals and incubated for 12 h; this modified induction procedure was only applied in strains that have defects in trap formation for detecting low trap formation. Nematodes were then removed by washing with ddH₂O. Trap numbers were quantified after 24 h using a Zeiss Stemi 305 microscope (40x magnification) with a Zeiss Axiocam ERc 5s camera. To evaluate the effect of glucose availability on trap suppression, LNM agar Petri plates were supplemented with a series of glucose concentrations prepared from a 2 M glucose stock solution. Sorbitol was used as an osmotic control to distinguish glucose-specific effects from changes in medium osmolarity. LNM agar Petri plates were supplemented with sorbitol at concentrations matching the corresponding glucose treatments. After applying glucose or sorbitol solutions to fungal cultures, plates were air-dried briefly and trap induction was assayed following the same procedure.

### Generation of gene deletion mutants and complemented strains

Gene deletions were performed in either wild type or *ku70* mutant backgrounds in *A. oligospora*. The *ku70* mutant strain was chosen for its deficiency in non-homologous end joining (NHEJ), which enhances homologous recombination efficiency [44]. Knockout cassettes comprised three fragments, amplified from the wild type using oligonucleotides listed in S27 Table: (1) 1.5 kb genomic sequence upstream of the coding region of the targeted gene, (2) a 1.5 kb genomic sequence identical to the downstream of the coding region, and (3) a nourseothricin acetyltransferase gene (*NAT1*) which had been amplified from vector pRS41N [53]. The 5′ end of fragment (2) and 3′ end of fragment (1) were designed to include 20 bp and 21 bp sequences, respectively, identical to sequences adjacent to the *NAT1* gene. These short homology sequences could later be used to fuse the resulting amplicons together by PCR. The three fragments were amplified individually and then assembled by PCR with primers targeting the terminal regions of the 5′ and 3′ UTRs. The assembled knockout constructs were introduced into protoplasts via PEG-mediated transformation as described below [9].

Protoplasts were obtained from conidia cultured in 50 mL PDB and incubated at 25°C with shaking at 200 rpm for 24 h. Hyphae were collected via centrifugation, washed with MN buffer (0.3 M MgSO₄, 0.3 M NaCl), and were digested overnight in 10 mL of 50 mg/mL VinoTastePro enzyme mix (Novonesis) dissolved in MN buffer at 25°C with shaking at 200 rpm. Protoplasts were filtered through Miracloth (EMD Millipore / Merck) and washed with STC buffer (1.2 M sorbitol, 50 mM CaCl₂, 10 mM Tris–HCl, pH 7.5). Next, approximately 3.5 × 10⁵ protoplasts were mixed with 2.5 μg of DNA targeting construct and incubated on ice for 30 minutes. Five volumes of PTC buffer (40% PEG 4000 [w/v]; 10 mM Tris–HCl, pH 7.5; 50 mM CaCl₂) were then added and incubated at room temperature for 20 minutes. The protoplast suspension was subsequently mixed with molten regeneration agar (1% agar, 0.5 M sucrose, 0.3% yeast extract, 0.3% casein acid hydrolysate) containing either 100 μg/mL hygromycin B (ANT-HG-5 InvivoGen) or 250 μg/mL clonNAT (N-500 GoldBio) and poured into Petri dishes. A total of six targeted gene deletion mutants were obtained. Putative mutants were screened by PCR using primers flanking the deletion cassette and then validated by Southern blotting [9].

Complementation constructs contained wild type copies of the target gene, including ∼2 kb upstream and ∼1 kb downstream flanking sequences, fused to a G418/geneticin resistance marker [54]. The constructs were introduced into protoplasts of targeted mutant strains. Transformants were selected on medium containing 240 μg/mL G418 (CAS 108321-42-2, Geneticin) and verified by genotyping. For the *CRE1^Δ292-335^*expression construct, two fragments were amplified from the *CRE1* sequence, excluding the conserved region encoding amino acids 292-335, and fused with the geneticin resistance marker. Additionally, to assess Cre1 functional complementation, the *AoCRE1* promoter sequence (1.5 kb upstream) was fused to the coding region of *N. crassa CRE1* (*NcCRE1*), which was retrieved from the *N. crassa* OR74A reference genome (NCBI: GCA_000182925.2).

### Generation of GFP fusion constructs

For construction of the Cre1-GFP fusion expression plasmid, the *CRE1* coding sequence together with 2 kb of its upstream promoter region was amplified by PCR using primers listed in S27 Table. PCR reactions were performed with Phusion High-Fidelity DNA Polymerase (NEB). The resulting amplicon was cloned into a GFP expression vector containing a geneticin resistance cassette, which was kindly provided by Dr. Reinhard Fischer. Cloning was performed using the In-Fusion HD PCR Cloning Kit (Clontech). Positive clones were verified by sequencing and subsequently used for transformation experiments.

For construction of the Smt1-GFP fusion expression plasmid, the *SMT1* coding sequence together with 2 kb of its upstream promoter region was amplified by PCR using primers listed in S27 Table and cloned into the GFP expression vector described above. Positive clones were verified by sequencing and transformed into the *smt1* mutant strain. Transformants were selected on medium containing G418 and verified by genotyping. The functionality of the Smt1-GFP fusion protein was assessed by trap quantification after exposure to *C. elegans*.

### Protein sequence and structure analyses

Protein sequences for *Neurospora crassa* Cre1 (NCBI: EAA32758), *Aspergillus nidulans* CreA (UniProt: Q01981), and *Trichoderma reesei* CreA (NCBI: AAB01677) were obtained from the NCBI database. Protein sequences for *Neurospora crassa* Gsl-9 (NCBI: EAA33884), *Aspergillus nidulans* SmtA (NCBI: XP_663292), *Aspergillus nidulans* SmtB (UniProt: A0A1U8QNM5), *Candida albicans* Mts1 (NCBI: KAL1581151), and *Cryptococcus neoformans* Smt1 (NCBI: XP_012052547) were also obtained from the NCBI database. Multiple sequence alignments of Cre1 and Smt1 homologs were performed using T-Coffee [55]. The predicted protein structures of *A. oligospora* Cre1 and Smt1 were generated using AlphaFold2 [56] and visualized using UCSF Chimera [57].

### Phenotypic characterization of fungal strains

Trap quantification assays involved incubating fungal strains on 3.5 cm LNM agar Petri plates for 2 days at 25°C. Subsequently, thirty *C. elegans* animals at the young adult stage were added onto the fungal plates and incubated for 6 h. The nematodes were then removed by washing with ddH₂O. After 24 h, three random images within a 0.5 cm radius were captured using a Zeiss Stemi 305 microscope with a Zeiss Axiocam ERc 5s camera at 40x magnification. Trap numbers were quantified using Fungal Feature Tracker [58].

Conidiation images were taken from fungal strains cultured on 5 cm LNM agar Petri plates for 5 days at 25°C. Images were captured using a Zeiss Stemi 305 microscope with a Zeiss Axiocam ERc 5s camera at 80× magnification.

Trap morphology imaging was conducted on fungal samples that had been exposed to *C. elegans* for 6 h on LNM agar Petri plates, washed with ddH2O to remove nematodes, and then incubated for additional 24 h. Samples were stained for 10 minutes with 3 μL of SCRI Renaissance 2200 (0.02%) in 15 mL ddH₂O, rinsed twice with ddH₂O, and imaged with an LSM980 Airyscan 2 confocal microscope using a Plan-Apochromat 63×/1.4 oil differential interference contrast objective (Zeiss). Images were acquired using Zeiss ZEN 3.3 (blue edition) software in multiplex mode (SR-4Y).

For Smt1-GFP localization imaging, fungal strains were cultured on LNM agar plates with or without exposure to *C. elegans* and stained with SR2200 as described above. GFP fluorescence was used to detect Smt1-GFP localization. For quantification of Smt1-GFP fluorescence distribution, fluorescence intensity was measured in ImageJ (NIH) [59] along the inner and outer sides of vegetative hyphal tips and developing trap hyphal tips. Segmented lines with a width of 1.9 µm were drawn from the hyphal tip along both sides of the cell. For vegetative hyphae, fluorescence was measured over a 10 µm length from the hyphal tip, whereas for developing trap cells, fluorescence was measured along the curved apical region of bending trap hyphae. The mean fluorescence intensity from each side was calculated, and Smt1-GFP distribution was expressed as the inner/outer fluorescence ratio. Statistical analysis was performed in GraphPad Prism 9 (GraphPad Software) using two-tailed unpaired Student’s t-test.

For filipin staining, fungal cultures on 5 cm LNM agar Petri plates were incubated for 3 days (*smt1* mutants for 7 days). For nematode-induced trap conditions, approximately 200 *C. elegans* were applied for 4h and removed by rinsing twice with ddH_2_O; control plates were processed in parallel without nematode exposure. A modified filipin staining procedure was used, based on previously published methods [60]. Filipin III (Cayman Chemical) was prepared as a 1 mg/mL stock solution in DMSO and diluted to a working concentration of 25 μg/mL with ddH_2_O. 50 μl of diluted filipin solution was applied directly to fungal cultures and incubated in the dark for 5 minutes. Cultures were rinsed twice with ddH₂O before examination by microscopy. Filipin fluorescence intensity profiles were quantified in ImageJ (NIH) [59] by measuring fluorescence along a rectangular region of interest (ROI; 5 µm length × 1.9 µm width) positioned starting at the hyphal tip and extending along the hyphae. For control intensity profiles, an ROI of identical settings was placed at a randomly selected region of vegetative hyphae within the same image (chosen away from trap and vegetative hyphal tips). Relative fluorescence intensity was calculated by subtracting the vegetative hyphae control profile from the hyphal tip profile. Each replicate corresponded to one hyphal tip from an independent trap, and the number of hyphal tips analyzed for each strain is indicated in the figure. Statistical analysis was performed in GraphPad Prism 9 (GraphPad Software) using a two-way mixed-effects model (REML) with strain and distance as fixed effects and repeated measurements across distance within each replicate. Pairwise analyses were performed between strains (WT vs *smt1*, *smt1* vs *smt1SMT1*, *cre1* vs *cre1CRE1*, and *cre1* vs *cre1P_GPD1_::SMT1*). Multiple comparisons were conducted within each distance point (row) by comparing each cell mean with every other cell mean in that row, with Šidák correction for multiple testing. Differences were considered statistically significant at P < 0.05.

### Protein Extraction and Western blot analysis

Culturing conditions and sample collection were performed as previously described [12]. Fungal cultures were cultured on 9 cm LNM agar Petri plates for 5 days at 25°C. The fungal cultures were collected using cell scrapers and immediately frozen in liquid nitrogen followed by lyophilization. For total protein extraction, samples were disrupted using 0.5 mm glass beads to grind the lyophilized hyphae into Mycelia powder. The powder was resuspended in a non-denaturing extraction buffer (20 mM Tris–HCl, pH 8.0, 137 mM NaCl, 10% glycerol, 1% Triton X-100, 1 mM phenylmethylsulfonyl fluoride (93482 sigma Aldrich), and 1× protease inhibitor cocktail (P2714, Sigma-Aldrich)). Protein concentration was determined using the Bradford protein assay (Bio-Rad) [61]. Proteins were separated on 4–12% gradient polyacrylamide gels (GenScript), transferred to polyvinylidene difluoride (PVDF) membranes and blocked in TBST buffer (20 mM Tris, 150 mM NaCl, and 0.1% Tween 20, pH 7.6) containing 1% (w/v) casein at room temperature for 2 h. Membranes were then hybridized with anti-GFP (Abcam, ab290) and anti-tubulin (Abcam, ab1849970), each diluted 1:1,000 in blocking solution, and incubated overnight at 4°C. After the membranes had been washed with TBST, they were incubated for 1 h at room temperature with a 1:8,000 dilution of AffiniPure Donkey anti-Rabbit IgG (Jackson ImmunoResearch, 715-035-152). For signal detection, membranes were incubated with enhanced chemiluminescence detection system (Bio-Rad Clarity Western ECL substrate) and subjected to standard film-developing procedure. Band intensities were quantified using ImageJ (NIH) [59], and target protein signals were normalized to the tubulin loading control.

### Total RNA isolation, RNA-seq library preparation, data analysis, and qPCR analysis

Total RNA isolation, RNA-seq library preparation, and data analysis were performed as previously described [9]*. A. oligospora* wild type and *cre1* mutant strains were cultured on 9 cm LNM agar Petri plates for 5 days and 7 days respectively at 25°C. Then, around 1000 young adult *C. elegans* were placed on each plate, and the nematodes were washed away after 4 h with ddH₂O. Each fungal sample was collected with a cell scraper and centrifuged at 4696 g, supernatant was removed, and the samples were frozen in liquid nitrogen, followed by lyophilization. Total RNA was extracted by the Trizol-Phenol-Chloroform method [62], and treated with Turbo DNase (Thermo Fisher Scientific), and cleaned by ethanol precipitation. RNA quality and quantity were determined using Bioanalyzer (Thermo Fisher Scientific) and Qubit systems (Thermo Fisher Scientific), respectively. The cDNA libraries for total RNA samples were prepared by the Genomics Core Facility (Institute of Molecular Biology, Academia Sinica), following standardized protocols [9]. Sequencing was performed on the Illumina NextSeq 500 platform for 150 cycles (high output) and 75 bp paired-end reads were obtained. Raw sequencing data were obtained in FASTQ format and subjected to quality control using FastQC to assess read quality and adapter contamination. High-quality reads were aligned to the reference fungal genome using STAR (Spliced Transcripts Alignment to a Reference) [63] with default parameters for paired-end RNA-seq reads. Gene-level quantification was performed using RSEM (RNA-Seq by Expectation-Maximization) [64], which calculated transcript abundances in terms of TPM (Transcripts Per Million) and raw read counts. The output was compiled into a gene expression matrix for downstream analysis.

To conduct differential gene expression analysis, sample groups were defined based on genotype and experimental conditions (*cre1* vs WT / WT+*C. elegans* vs WT / *cre1*+*C. elegans* vs *cre1*). Pairwise comparisons between experimental conditions were used to identify differentially expressed genes (DEGs). Genes with an adjusted P-value (FDR) < 0.05 and a beta value threshold of ≥1.5 or ≤ -1.5 were considered significant DEGs, using Sleuth 0.29.0 [65]. Results were visualized using volcano plots to highlight DEG distribution. To identify *cre1*-dependent nematode responsive genes related to trap formation, transcript clusters were obtained using Clust [66] based on TPM values across all conditions and replicates. For clusters C7 and C10, genes were filtered based on the wild type response (WT + *C. elegans* vs WT) using thresholds of FDR-adjusted P-value < 1×10⁻⁵ and |beta| ≥ 1.5. Remaining DEGs were run through Omicsbox 2.0.36 [67]. For functional enrichment analysis, a *P*-value < 0.01 (Fisher’s exact test) was used as the significance threshold. The initial results were obtained for “Interpro GO IDs”. Heat maps were built on GraphPad Prism (GraphPad Software, LLC, San Diego, CA, USA). Principal component analysis results are provided in S9 Table. DEG lists for the three pairwise comparisons are provided in S10-S12 Tables. The full TPM expression dataset used for clustering and the resulting cluster assignments are provided in S13 Table. The C7-refined and C10-refined gene sets are provided in S14 Table, and the corresponding GO enrichment analysis is provided in S15 Table.

Quantitative PCR was performed using the QuantStudio 12K Flex Real-Time PCR System (Applied Biosystems, Thermo Fisher Scientific). Each 20 μL reaction contained cDNA template, 100 nM of each primer, and 10 μL of Fast SYBR Green Master Mix (Applied Biosystems, Thermo Fisher Scientific). The cycling program consisted of a 95 °C denaturation step for 20 s, followed by 40 cycles of 95 °C for 1 s and 60 °C for 20 s. A final melt-curve step was included to verify amplification specificity. At least three biological replicates were analyzed for each condition. CT values were normalized to the reference gene *GPD1*, and relative quantification was calculated using the 2^−ΔCT^ method. For analysis of the *CRE1*^Δ292–335^ transcript, the same primer pair used for endogenous *CRE1* expression analysis was used.

## Acknowledgments

We thank Sue-Ping Lee, Yae-Huei Liou, and Chun-Yung Chang from the Imaging Core for their assistance with imaging, Shu-Yun Tung from the Genomics Core for her help with Illumina sequencing, and Chen-Hsin Yu and Hsin-Nan Lin from the Bioinformatics Core for their assistance with transcriptomic profile analyses at IMB, Academia Sinica. We also thank Drs. Yi-Fang Tsay, Hsou-Min Li, Jun-Yi Leu, and Rey-Huei Chen from IMB, Academia Sinica, for their comments and suggestions on the manuscript.

## Supporting information

S1 Table. *Arthrobotrys oligospora* strains used in this study.

S2 Table. Trap Number induced by *C. elegans* in wild type *A. oligospora* on LNM plates with different concentrations of glucose or sorbitol.

S3 Table. Trap numbers induced by *C. elegans* in wild type, *cre1* and *cre1CRE1* strains on LNM or LNM supplemented with 150mM glucose or sorbitol.

S4 Table. Trap numbers induced by *C. elegans* in the WT, *cre1*, and *cre1CRE1-GFP* strains.

**S5 Table**. Quantification of gene expression analysis of *CRE1* in the wild type strain on LNM agar media with or without glucose supplementation and with or without exposure to *C. elegans*.

S6 Table. Quantification of Cre1 abundance in the Cre1-GFP strain with or without glucose supplementation and to *C. elegans* exposure.

S7 Table. Quantification of the trap numbers induced by *C. elegans* in WT, cre1, *cre1CRE1*^Δ292-335^, *cre1Nc-CRE1, and cre1CRE1* grown on LNM.

S8 Table. Quantitative gene expression analysis of *CRE1* transcripts in wild type, *cre1*, and *cre1CRE1*^Δ292–335^ strains grown on LNM agar media with or without exposure to *C. elegans*.

S9 Table. Principal component analysis of *A. oligospora* transcriptomes from wild type (WT) and *cre1* mutant strains with or without exposure to *C. elegans*.

S10 Table. Differential gene expression (DEG) analysis in the *cre1* mutant relative to wild type.

S11 Table. Differential gene expression (DEG) analysis in the wild type with or without *C. elegans* exposure.

S12 Table. Differential gene expression (DEG) analysis in the *cre1* mutant with or without *C. elegans* exposure.

S13 Table. Cluster analysis of transcript expression profiles in the wild type and *cre1* mutant with or without *C. elegans* exposure.

**S14 Table. DEGs from Cluster C7-refined and C10-refined datasets.**

S15 Table. GO enrichment analysis of C7-refined and C10-refined dataset. S16 Table. Quantification of the trap numbers induced by *C. elegans* in the wild type and in strains overexpressing selected Cluster C7-refined genes.

S17 Table. Quantification of the trap numbers induced by *C. elegans* in the wild type and the indicated gene deletion strains.

S18 Table. Quantification of trap numbers formed on LNM in the WT, *smt1*, *smt1SMT1*, *cre1*, *cre1P_GPD1_::SMT1*, *ste12*, and *ste12P_GPD1_::SMT1* strains after exposure to *C. elegans*.

S19 Table. Quantitative gene expression analysis of *SMT1* gene expression in the wild type, *cre1*, *cre1P_GPD1_::STE12*, *ste12*, and *ste12P_GPD1_::CRE1* strains.

S20 Table. *STE12* gene expression in the wild type and *cre1* mutant strains with or without *C. elegans*.

S21 Table. *CRE1* gene expression in the wild type and *ste12* mutant strains with or without *C. elegans*.

S22 Table. Quantitative gene expression analysis of *STE12* in the *cre1* mutant strain cultured on PDA plate.

S23 Table. Quantitative gene expression analysis of *CRE1* in the *ste12* mutant strain culture on PDA plate.

S24 Table. Quantification of the trap numbers induced by *C. elegans* in the WT, *smt1*, and *smt1SMT1-GFP* strains.

S25 Table. Relative Smt1-GFP fluorescence intensity ratios measured between the left and right sides of vegetative hyphae (VH) before *C. elegans* exposure and between the inner and outer sides of trap hyphae (TH) 4 h after *C. elegans* exposure.

S26 Table. Relative filipin fluorescence intensity profiles measured along the apical region of trap and vegetative hyphal tips of WT, *smt1*, *smt1SMT1*, *cre1*, *cre1P_GPD1_::SMT1*, and *cre1CRE1* strains on LNM with or without *C. elegans* exposure.

S27 Table. Primer sequences used in the experiments.

---

**S1 Fig.**
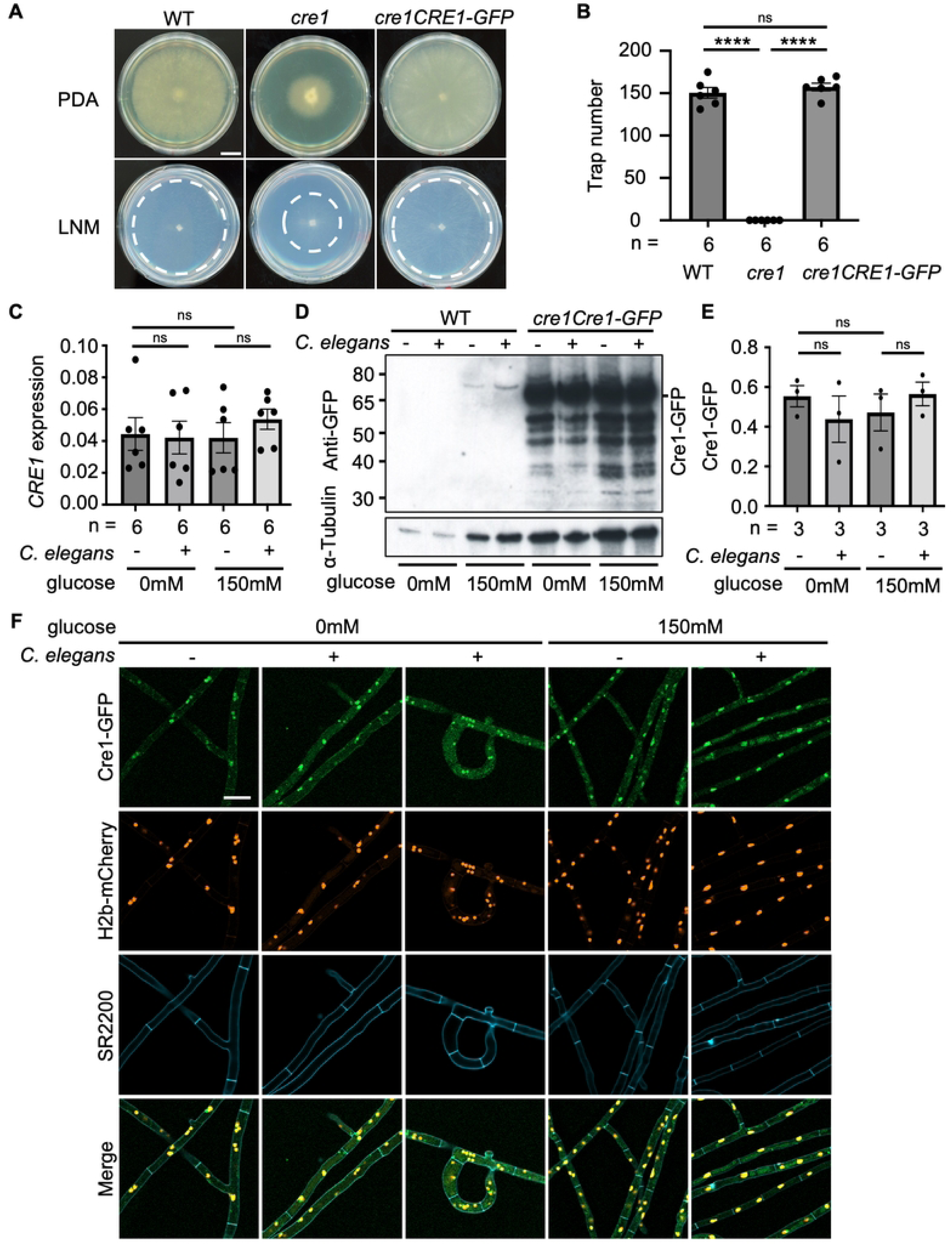
*CRE1* expression is stable and remains in nucleus in response to *C. elegans.* (A) Representative images of colony morphology of wild type, *cre1*, and *cre1CRE1-GFP* strain cultured on PDA or LNM agar Petri plates (scale bar = 1 cm). (B) Quantification of the trap numbers induced by *C. elegans* in the WT, *cre1*, and *cre1CRE1-GFP* strain. Statistical comparisons were performed among the strains. (data represent mean ± SEM; asterisks represent significance levels of two-tailed unpaired Student’s t-test; ****P< 0.0001, ns: Not significant). WT, wild type. (C) Quantitative gene expression analysis of *CRE1* in the wild type strain on LNM agar media with or without glucose supplementation and with or without exposure to *C. elegans*. *GPD1* was used as a normalization control. All comparisons were not statistically significant (ns) by two-tailed unpaired Student’s t test; therefore, no asterisks are shown. (data represent mean ± SEM) (D) Protein expression profile of Cre1 in the wild type and *cre1Cre1-GF*P strains with or without glucose supplementation and to *C. elegans* exposure. Total protein was extracted from each sample, with WT used as a negative control. Cre1-GFP (74.4 kDa) was detected by western blotting using anti-GFP antibody. Tubulin was detected using an anti-tubulin antibody. (E) Quantification of Cre1 abundance in the Cre1-GFP strain with or without glucose supplementation and to *C. elegans* exposure. Tubulin detected by anti-tubulin antibody was used to normalize Cre1– GFP signal intensity. (F) Subcellular localization of Cre1-GFP in the absence or presence of *C. elegans on LNM agar and on LNM agar supplemented with glucose*. GFP fluorescence (green) indicates Cre1 localization, the histone marker H2B-mCherry (red) indicates nuclei. SR2200 (blue) indicates cell walls. Traps can be seen in the third column.

**S2 Fig.**
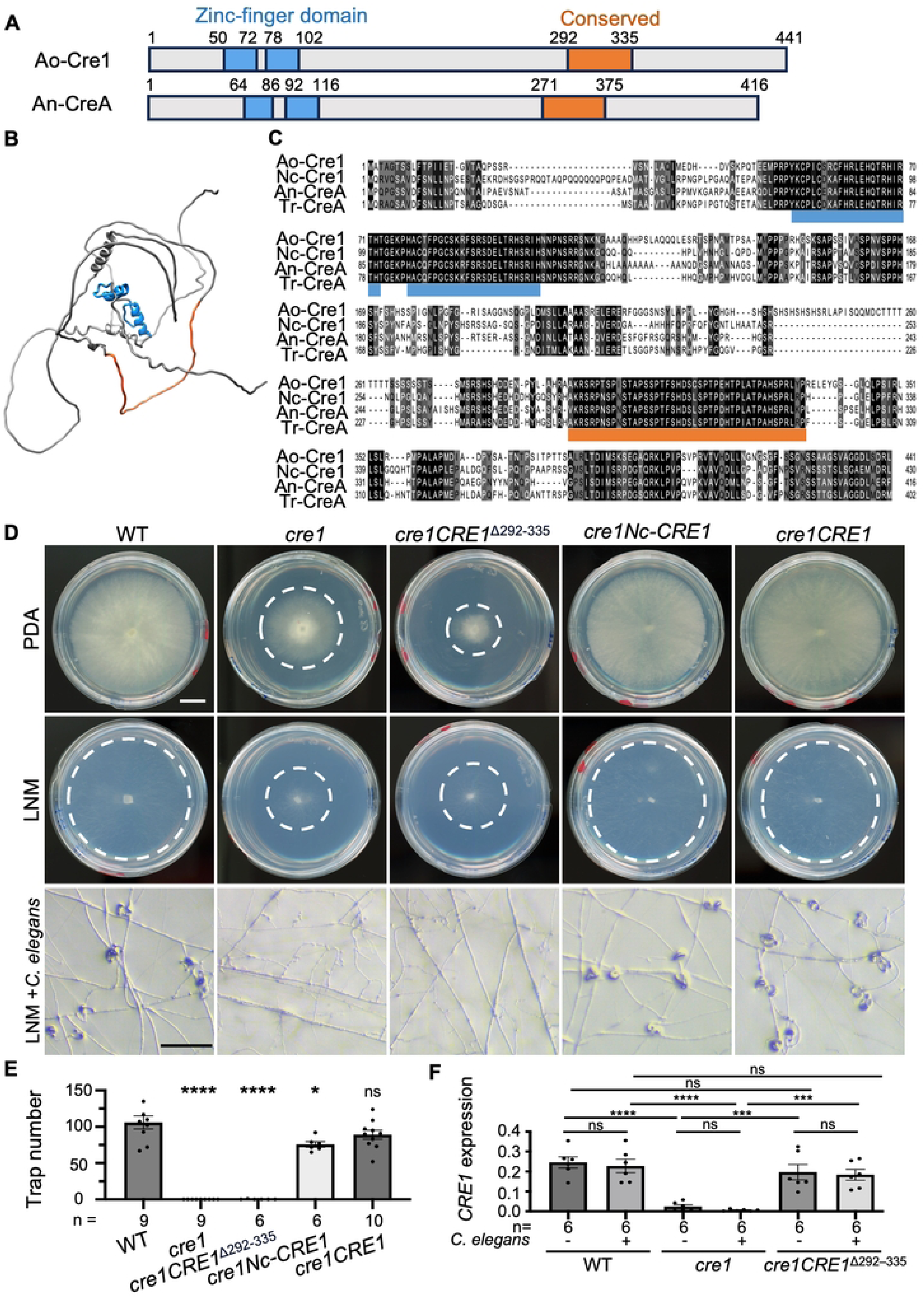
Cre1 292-335 region is highly conserved across filamentous fungi and is important to its function in *A. oligospora*. (A) Domain organization of Cre1 from *A. oligospora* and *A. nidulans*, highlighting zinc finger DNA-binding domains (blue) and conserved region (orange) specific to *A. nidulans*. (B) Predicted protein structure of *A. oligospora* Cre1. The zinc-finger DNA-binding domain and the conserved region spanning residues 292-335 are highlighted in blue and orange respectively. (C) Protein sequence alignment of Cre1 from *A. oligospora* (Ao), *N. crassa* (Nc), *T. reesei* (Tr), and *A. nidulans* (An). Different levels of shading indicate degrees of sequence similarity, with higher intensity shown in darker black. (D) Colony morphology and trap images of wild type, *cre1*, *cre1CRE1*^Δ292-335^, *cre1Nc-CRE1*, and *cre1CRE1* strains in *A. oligospora* cultured on PDA or LNM (Scale bar = 200 μm). Representative brightfield images show trap induction by *C. elegans* in the same strains (Scale bar = 200 μm). (E) Histogram shows quantification of the trap numbers induced by *C. elegans* in strains grown on LNM. (F) Quantitative gene expression analysis of *CRE1* transcripts in wild type, *cre1*, and *cre1CRE1*^Δ292–335^ strains grown on LNM agar media with or without exposure to *C. elegans*. *GPD1* was used as a normalization control. (data represent mean ± SEM; asterisks represent significance levels according to two-tailed unpaired Student’s t-test; *P<0.05, ***P<0.001, ****P< 0.0001, ns: Not significant).

**S3 Fig.**
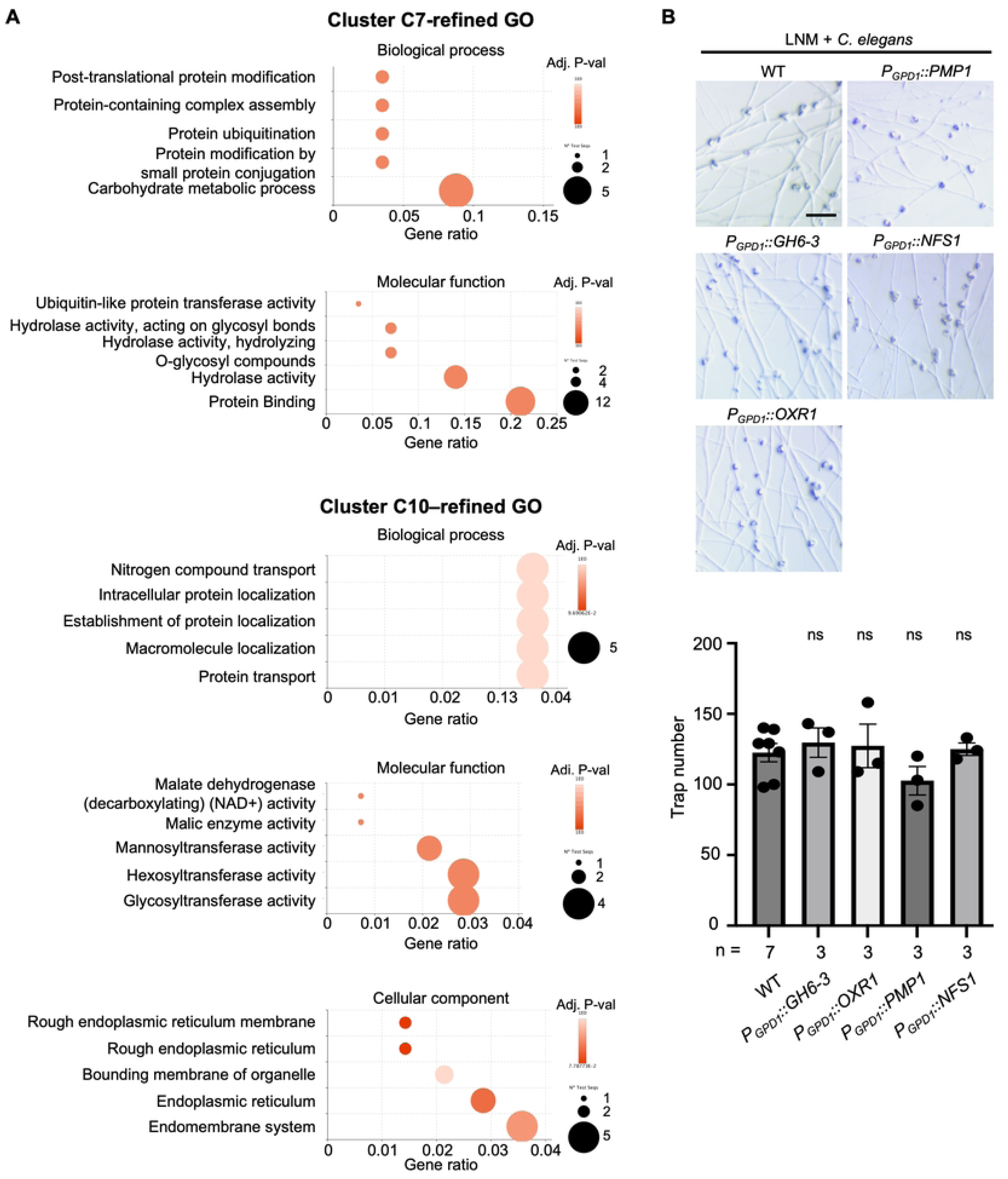
Gene Ontology analyses of Cluster C7-refined and Cluster C10-refined genes. (A) Gene Ontology (GO) enrichment analysis of DEGs in C7**-**refined and C10**-**refined gene sets. In the bubble plots, the gene ratio represents the number of genes annotated with a given GO term divided by the total number of genes in the respective refined set. Dot size reflects the number of genes associated with each GO term, and dot color corresponds to enrichment significance (-log_10_P-value, Fisher’s exact test). X-axis, labeled “Gene ratio”, indicates the proportion of genes in the set annotated with the indicated term. Y-axis labels indicate GO term names. The Gene ratios and adjusted P-values indicated that no terms were identified by GO term analysis as strong candidates to explain the function of Cre1 in trap formation. (B) Representative brightfield images of the traps induced by *C. elegan*s. Images provide comparisons between the wild type strain and strains overexpressing selected genes from Cluster C7-refined (Scale bar = 200 μm). Histogram shows quantification of the trap numbers induced by *C. elegans* in the wild type and in strains overexpressing selected Cluster C7-refined genes. (data represent mean ± SEM; asterisks represent significance levels according to two-tailed unpaired Student’s t-test; ns: Not significant).

**S4 Fig.**
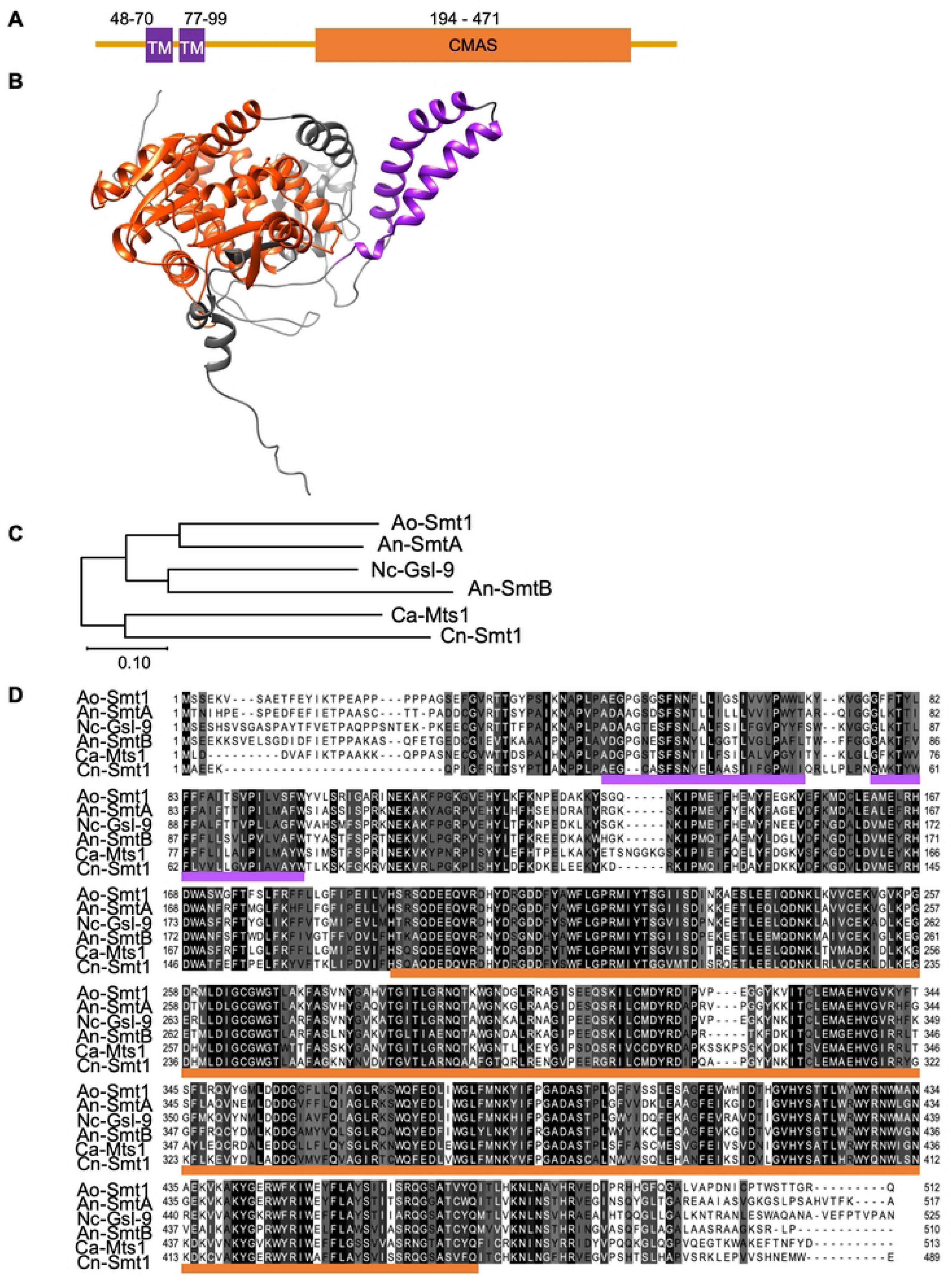
Smt1 is highly conserved across fungi and maintains a characteristic transmembrane-CMAS domain structure. (A) Domain organization of Smt1 proteins in *A. oligospora*, showing the predicted transmembrane (TM) regions and the CMAS-family methyltransferase domain. (B) Predicted protein structure of *A. oligospora* Smt1. The transmembrane domain is highlighted in purple and the CMAS region is highlighted in orange. (C) Phylogenetic tree of Smt1 homologs in fungi constructed using the maximum-likelihood method in MEGA11. Species abbreviations: *A. oligospora* (Ao), *A. nidulans* (An), *N. crassa* (Nc), *C. albicans* (Ca), and *C. neoformans* (Cn). (D) Protein sequence alignment of Smt1 from *A. oligospora* and orthologs from model fungi. Species abbreviations: *A. oligospora* (Ao), *A. nidulans* (An), *N. crassa* (Nc), *C. albicans* (Ca), and *C. neoformans* (Cn). Different levels of shading indicate degrees of sequence similarity, with higher intensity shown in darker black. Transmembrane domains are underlined in purple and the CMAS domain is underlined in orange.

**S5 Fig.**
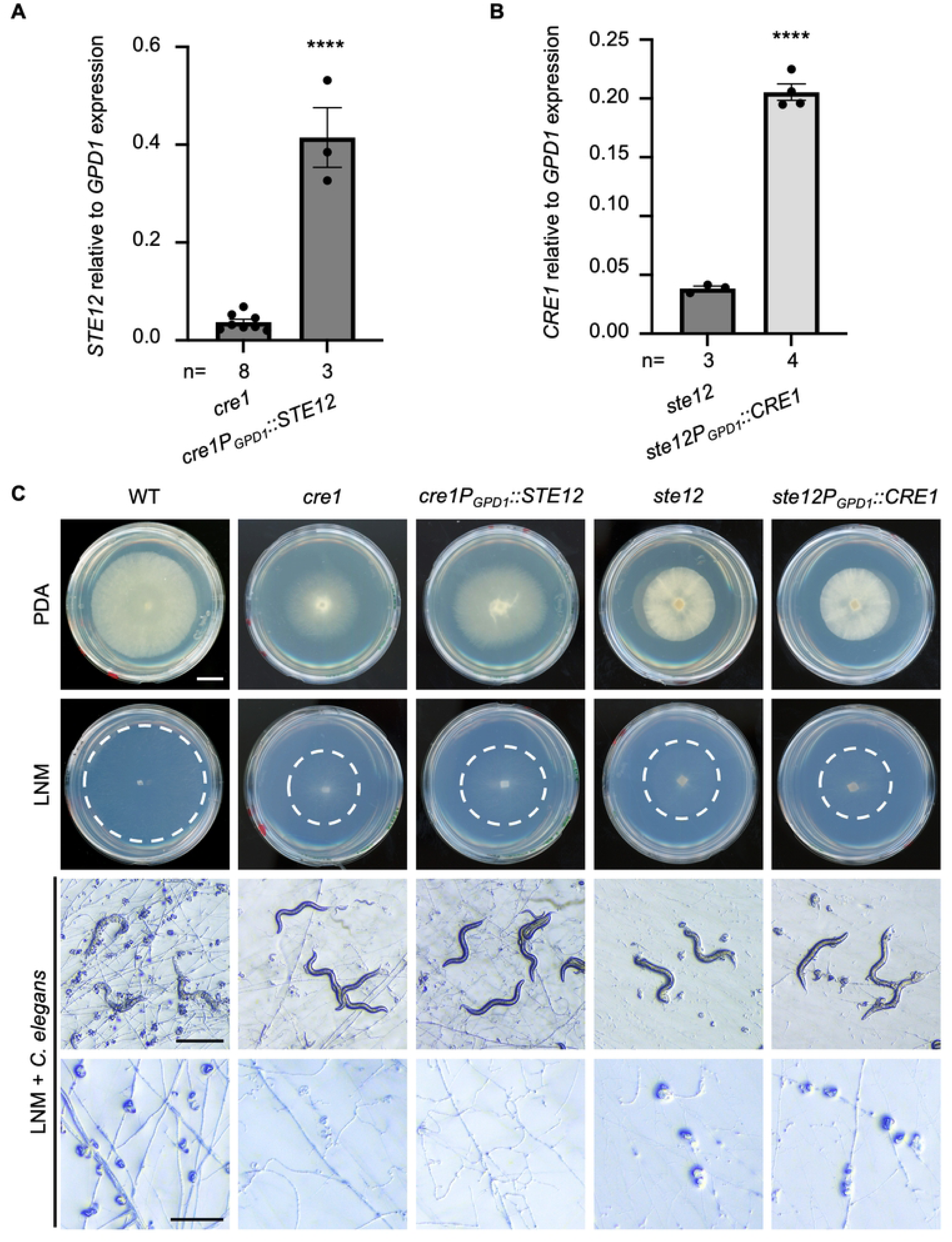
Trap quantification in wild type and mutant strains induced by *C. elegans*. (A) Representative images of colony morphology of wild type, *smt1*, *smt1SMT1*, *cre1*, *ste12*, *cre1P_GPD1_::SMT1*, and *ste12P_GPD1_::SMT1* cultured on LNM agar Petri plates (scale bar = 1 cm). (B) Quantification of trap numbers formed on LNM in the WT, *smt1*, *smt1SMT1*, *cre1*, *cre1P_GPD1_::SMT1*, *ste12*, and *ste12P_GPD1_::SMT1* strains after exposure to *C. elegans*. Statistical comparisons were performed among the strains. Data represent mean ± SEM; asterisks represent significance levels according to two-tailed unpaired Student’s t-test; ***P<0.001, ****P< 0.0001, ns: Not significant). WT, wild type.

**S6 Fig.**
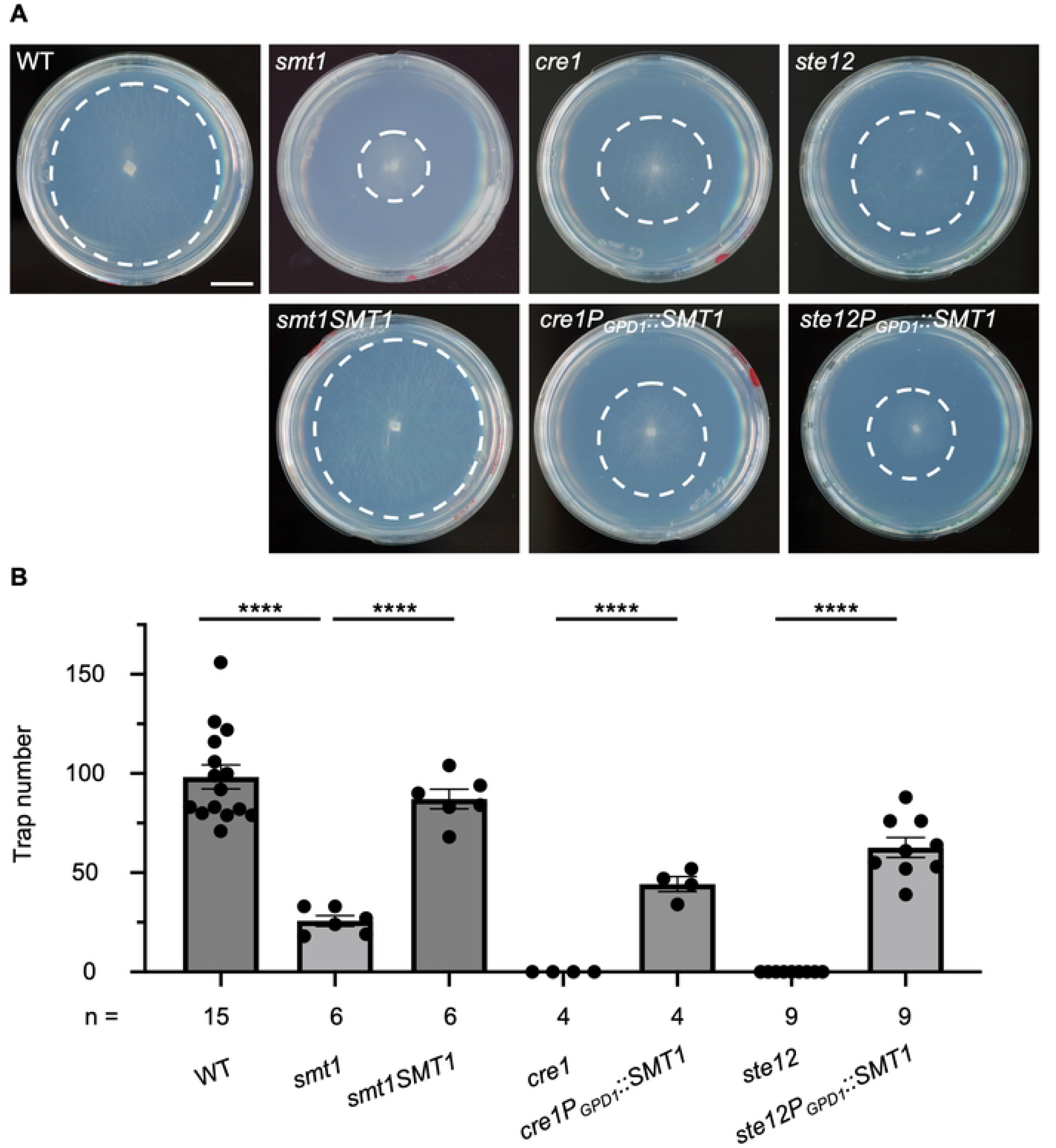
Reciprocal overexpression of *STE12* and *CRE1* fails to restore trap formation in *cre1* and *ste12* mutants. (A) Quantitative gene expression analysis of *STE12* in the *cre1* mutant strain cultured on PDA plate. (B) Quantitative gene expression analysis of *CRE1* in the *ste12* mutant strain culture on PDA plate. *GPD1* was used as a normalization control. (data represent mean ± SEM; asterisks represent significance levels of two-tailed unpaired Student’s t-test; ****P< 0.0001). (C) Representative images of colony morphology of WT, *cre1*, *cre1P_GPD1_::STE12*, *ste12*, and *ste12P_GPD1_::CRE1* strains cultured on PDA or LNM agar Petri plates (Scale bar = 1 cm). Brightfield images show strains grown on LNM agar and exposed to *C. elegans* (scale bar = 500 μm), and higher magnification images of traps used for trap quantification after nematodes were removed (Scale bar = 200 µm). WT, wild type.

**S7 Fig.**
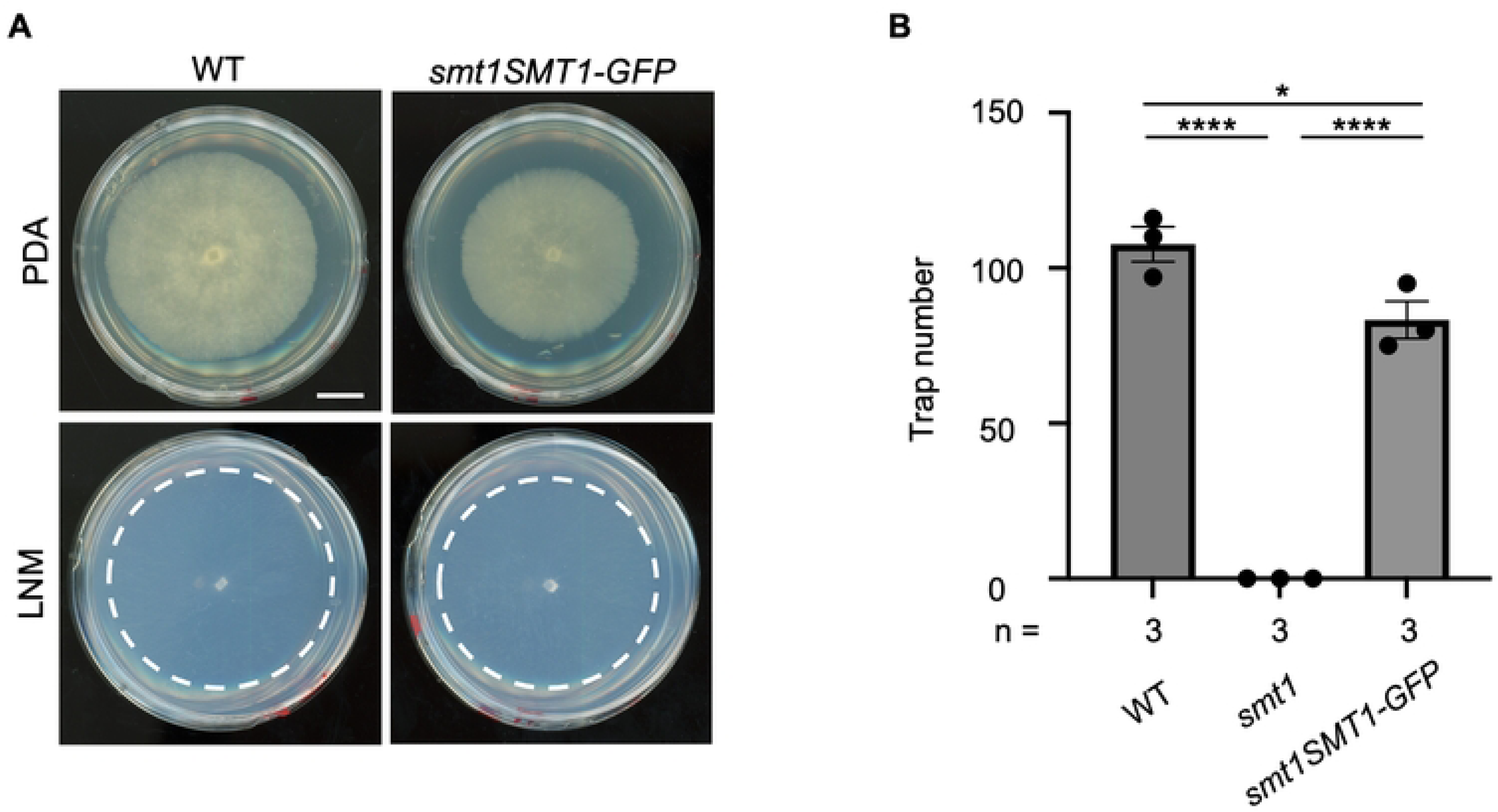
Smt1-GFP is functional and restores growth and trap formation in *smt1* mutant. (A) Representative images of colony morphology of WT and *smt1SMT1-GFP* strains cultured on PDA or LNM agar Petri plates (Scale bar = 1 cm). (B) Quantification of the trap numbers induced by *C. elegans* in the WT, *smt1*, and *smt1SMT1-GFP* strains. Statistical comparisons were performed among the strains. (data represent mean ± SEM; asterisks represent significance levels of two-tailed unpaired Student’s t-test; *P<0.05, ****P< 0.0001). WT, wild type.

